# DREAM represses distinct targets by cooperating with different THAP domain proteins

**DOI:** 10.1101/2020.08.13.249698

**Authors:** Csenge Gal, Francesco Nicola Carelli, Alex Appert, Chiara Cerrato, Ni Huang, Yan Dong, Jane Murphy, Julie Ahringer

**Author notes:** Contributed equally (Csenge Gal, Francesco Nicola Carelli).

## Abstract

The DREAM (DP, Retinoblastoma [Rb]-like, E2F, and MuvB) complex controls cellular quiescence by repressing cell cycle and other genes, but its mechanism of action is unclear. Here we demonstrate that two C. elegans THAP domain proteins, LIN-15B and LIN-36, co-localize with DREAM and function by different mechanisms for repression of distinct sets of targets. LIN-36 represses classical cell cycle targets by promoting DREAM binding and gene body enrichment of H2A.Z, and we find that DREAM subunit EFL-1/E2F is specific for LIN-36 targets. In contrast, LIN-15B represses germline specific targets in the soma by facilitating H3K9me2 promoter marking. We further find that LIN-36 and LIN-15B differently regulate DREAM binding. In humans, THAP proteins have been implicated in cell cycle regulation by poorly understood mechanisms. We propose that THAP domain proteins are key mediators of Rb/DREAM function.

## INTRODUCTION

During animal development, cell proliferation is tightly controlled and differentiated cells spend the majority of the time in a quiescent, non-dividing, state. The regulation of quiescence is crucial, as uncontrolled proliferation can lead to tumour formation, whilst premature senescence is associated with ageing. Despite its importance, mechanisms of quiescence regulation remain poorly understood.

The Retinoblastoma family of pocket proteins (Rb, p130, p107) are key regulators of the cell division cycle, regulating progression from G_1_-S phase and maintaining the G_0_ state via transcriptional repression of proliferation promoting genes (Dick and Rubin, 2013). The majority of cancers disable Rb protein function or alter its regulation (Liu et al., 2013; Rashid et al., 2011; Sadasivam and DeCaprio, 2013). Loss of Rb also leads to developmental defects (Du et al., 1996; Lee et al., 1992; Lu and Robert Horvitz, 1998). A mechanistic understanding of Rb proteins is essential for understanding their roles in normal development and in cancerous transformations.

Of the Rb family of proteins p130 is the most highly expressed during stable cell cycle arrest, such as quiescence and senescence, through which it represses proliferation-promoting genes as part of a repressive complex called DREAM (Lewis et al., 2004; Litovchick et al., 2007) Litovchick et al., 2007, 2011; Schmit et al., 2007). In different organisms, disruption of DREAM leads to developmental defects, an increase in genomic instability, tumorigenesis and lethality (Hauser et al., 2012; Malumbres and Barbacid, 2009; Reichert et al., 2010; Schade et al., 2019). The mechanisms by which DREAM functions in these different processes is unclear.

The DREAM complex is highly conserved in subunit composition and function in animals (Sadasivam and DeCaprio, 2013). Mammalian DREAM is composed of an Rb-like protein p130 (or p107 in the absence of p130), an E2F (E2F4/E2F5), a dimerization partner (DP) protein, and MuvB proteins (LIN9, LIN54, LIN52, LIN37, RBBP4) (Litovchick et al., 2007; Schmit et al., 2007). As in mammals, C. elegans DREAM (LIN-35/Rb, DPL-1/DP, EFL-1/E2F, LIN-9, LIN-37, LIN-53, LIN-54 and LIN-52) represses cell cycle specific genes and others, including germline genes in somatic tissues (Goetsch et al., 2017; Korenjak et al., 2004; Latorre et al., 2015; Rechtsteiner et al., 2019). Since DREAM itself contains no known enzymatic activity, it is thought to repress targets through effector proteins. Indeed, such a role has been proposed for the Sin3B-HDAC complex in mammalian cells (Bainor et al., 2018; Rayman et al., 2002). In addition, we previously showed that repression of a subset of C. elegans DREAM targets involves deposition of HTZ-1/H2A.Z on their gene bodies (Latorre et al., 2015). To further mechanistic understanding, we undertook an RNAi screen for additional factors needed for repression of a DREAM target. Here we show that two THAP domain proteins function with DREAM by different mechanisms to repress distinct sets of targets.

## RESULTS

### An RNAi screen identifies novel regulators of Rb/DREAM targets

To identify proteins involved in DREAM transcriptional repression, we constructed a DREAM regulated reporter gene by fusing the promoter of the target sep-1 to a histone-eGFP coding region, and then carried out an RNAi screen for genes needed for reporter repression (Figure 1A). The screen was carried out in quiescent starved L1 larvae, which contain 550 non-dividing somatic cells and 2 germ cells. In wildtype starved L1s the P-sep-1::his-58::eGFP transgene is expressed in the germline and largely repressed in the soma (Figure 1B). In lin-35/Rb mutants, reporter expression is increased in the soma compared to the wildtype (Figure 1B). The RNAi screen targeted 1104 genes encoding nuclear proteins to identify genes that are required to prevent somatic expression of the P-sep-1::his-58::eGFP reporter (see Methods). Following RNAi knockdown, eGFP expression was measured using a worm sorter, which identified 36 genes for which knockdown caused reporter de-repression (Table S1), including seven out of eight DREAM components (lin-35/Rb, efl-1, dpl-1, lin-54, lin-9, lin-37 and lin-53), validating the screen. Others include components of the MCM complex, a number of RNA binding proteins, proteins required for kinetochore function and lin-36, which encodes a THAP domain containing protein.

**Figure 1.**
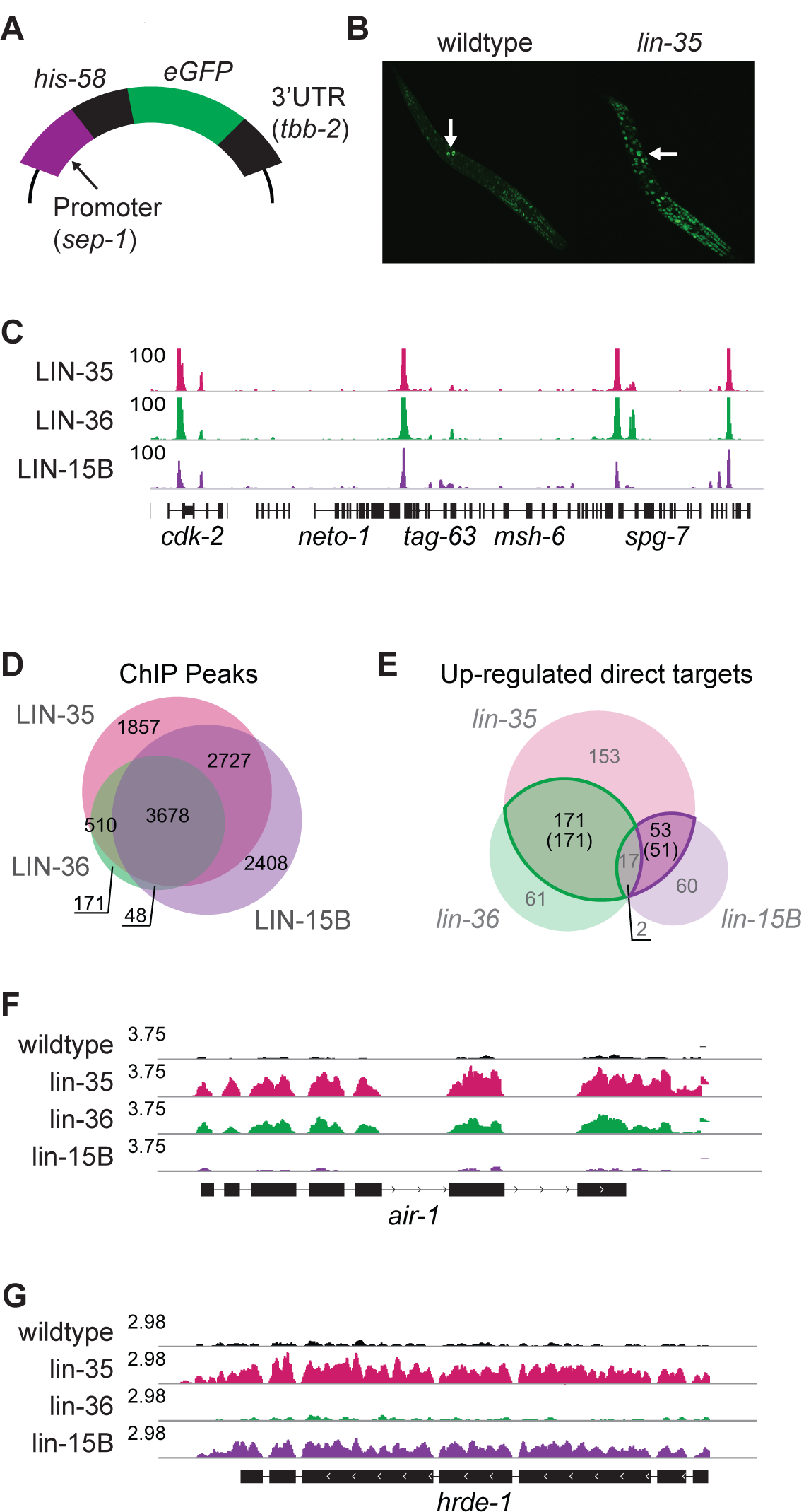
THAP-domain proteins LIN-36 and LIN-15B regulate Rb/DREAM targets. (A) p-sep-1::eGFP DREAM target reporter gene used for the RNAi screen. (B) lin-35 mutant animals have increased expression the p-sep-1::eGFP reporter relative to wild-type. Arrows indicate the two germ cells in starved L1 animals. (C) IGV view of linear BEADS normalised ChIP sequencing data for the indicated factors. (D) Overlap of ChIP peaks called for the indicated factors. (E) Overlap between direct targets in the indicated mutants. Number in parentheses indicate LIN-36-shared targets (green) and LIN-15B-shared targets (purple) (see Methods). (F) and (G) IGV view of RNA sequencing data of (E) a LIN-36-shared and (F) a LIN-15B-shared target.

LIN-36 was of particular interest, as its loss has been shown to cause cell cycle defects similar to those of DREAM mutants (Boxem and Van den Heuvel, 2002), but it has not been well characterized. LIN-36 contains a THAP domain, which is an atypical zinc finger DNA binding domain derived from a transposase (Clouaire et al., 2005; Roussigne et al., 2003). C. elegans has seventeen THAP or THAP-like domain-containing proteins, of which seven have been shown to genetically interact with lin-35/Rb (Table S1)(Boxem and Van den Heuvel, 2002; Ceron et al., 2007; Chesney et al., 2006; Ouellet and Roy, 2007; Poulin et al., 2005; Reddy and Villeneuve, 2004; Saito et al., 2004), suggesting a broad relationship between THAP domain proteins and LIN-35/Rb. Humans have 12 THAP domain proteins, THAP0 to THAP11, which have been implicated in diverse cellular processes, including the regulation of cell cycle genes (Cayrol et al., 2007; Ceron et al., 2007). Disruption of THAP proteins has also been linked to various diseases, including cancers (Balakrishnan et al., 2009; Gervais et al., 2013; Richter et al., 2017). We used RNAi to test whether other THAP domain genes are required for repression of the P-sep-1::his-58::eGFP reporter and found that LIN-15B is also needed (Table S1). Previous work showed that LIN-15B and LIN-35 share some transcriptional targets (Rechtsteiner et al., 2019), and LIN-15B has been implicated in negative regulation of the G_1_/S transition of the cell cycle (Boxem and Van den Heuvel, 2002). Here we investigate the roles of LIN-36 and LIN-15B in the repression of DREAM targets.

### LIN-36 and LIN-15B co-localize with LIN-35

To explore the relationship between LIN-35, LIN-36 and LIN-15B, we first compared their genome-wide binding patterns using ChIP-seq in wildtype starved L1 animals using antibodies to LIN-35 and LIN-15B and detecting LIN-36 by an endogenous GFP-tag (see Methods). We found that LIN-36 and LIN-15B both show a high degree of overlap with LIN-35, with 95% of LIN-36 and 72% of LIN-15B peaks overlapping a LIN-35 peak (Figures 1C, D, S1A, and Table S2). For each factor, most (59-69%) peaks overlap a promoter or enhancer with much of the remainder localising to repetitive elements (Figure S1B). Many of the repeat regions are marked by H3K9me2, supporting a possible connection between H3K9me2 and DREAM (Figure S1C; Rechtsteiner et al., 2019).

### LIN-36 and LIN-15B repress discrete sets of LIN-35 targets

We next compared the effects of loss of LIN-35, LIN-36 and LIN-15B on gene expression (Table S3). We used available null alleles lin-35(n745) and lin-15B(n744) and generated full deletion allele lin-36(we36) using CRISPR/Cas9 gene editing (see Methods). We also profiled the partial loss-of-function allele lin-36(n766). For all mutants, we observed that the primary effect was loss of repression (Table S3), and hence focused our work on direct repressed targets, which are defined as genes upregulated in lin-35, lin-36, or lin-15B mutants and bound by the corresponding factor (see Methods).

We observed that repressed targets of LIN-36 or LIN-15B each significantly overlap LIN-35/Rb targets (>21-fold enrichment, hypergeometric test P < 10^−76^), but strikingly, genes regulated by LIN-36 and LIN-15B are mostly distinct (Figure 1E, F). Here, we focus on genes directly regulated by LIN-35 and LIN-36 (LIN-36-shared targets; n=171) or regulated by LIN-35 and LIN-15B (LIN-15B-shared targets; n=51) (Table S3). Using gene ontology (GO) analyses, we found that LIN-36-shared targets are highly enriched for cell cycle and cell division terms (Table S3). No enriched GO terms were found for LIN-15B-shared targets (Table S3), however we observed that they have high germline expression specificity (Figure S2A, B; Table S3). LIN-36-shared targets and LIN-15B-shared targets also dramatically differ in the binding profiles of LIN-35, LIN-36 and LIN-15B, with higher signal for all three factors at LIN-36-shared targets compared to LIN-15B-shared targets (Figure S2C, D). Altogether, these observations suggest that LIN-15B-shared and LIN-36-shared genes represent two distinct classes of DREAM targets with potentially different regulation and functional roles.

### LIN-36 maintains gene body HTZ-1

We previously showed that transcriptional repression of a subset of DREAM target genes involves LIN-35-dependent enrichment of the histone variant H2A.Z/HTZ-1 over their gene bodies (gbHTZ-1) (Latorre et al., 2015). To assess whether LIN-36 and/or LIN-15B act with LIN-35 in facilitating gbHTZ-1, we first asked whether gene body enrichment of HTZ-1 was associated with either set of shared targets. Indeed, we observed that LIN-36-shared targets were more enriched for high gbHTZ-1 than LIN-15B-shared targets (Figures 2A, S3A; Table S4).

**Figure 2.**
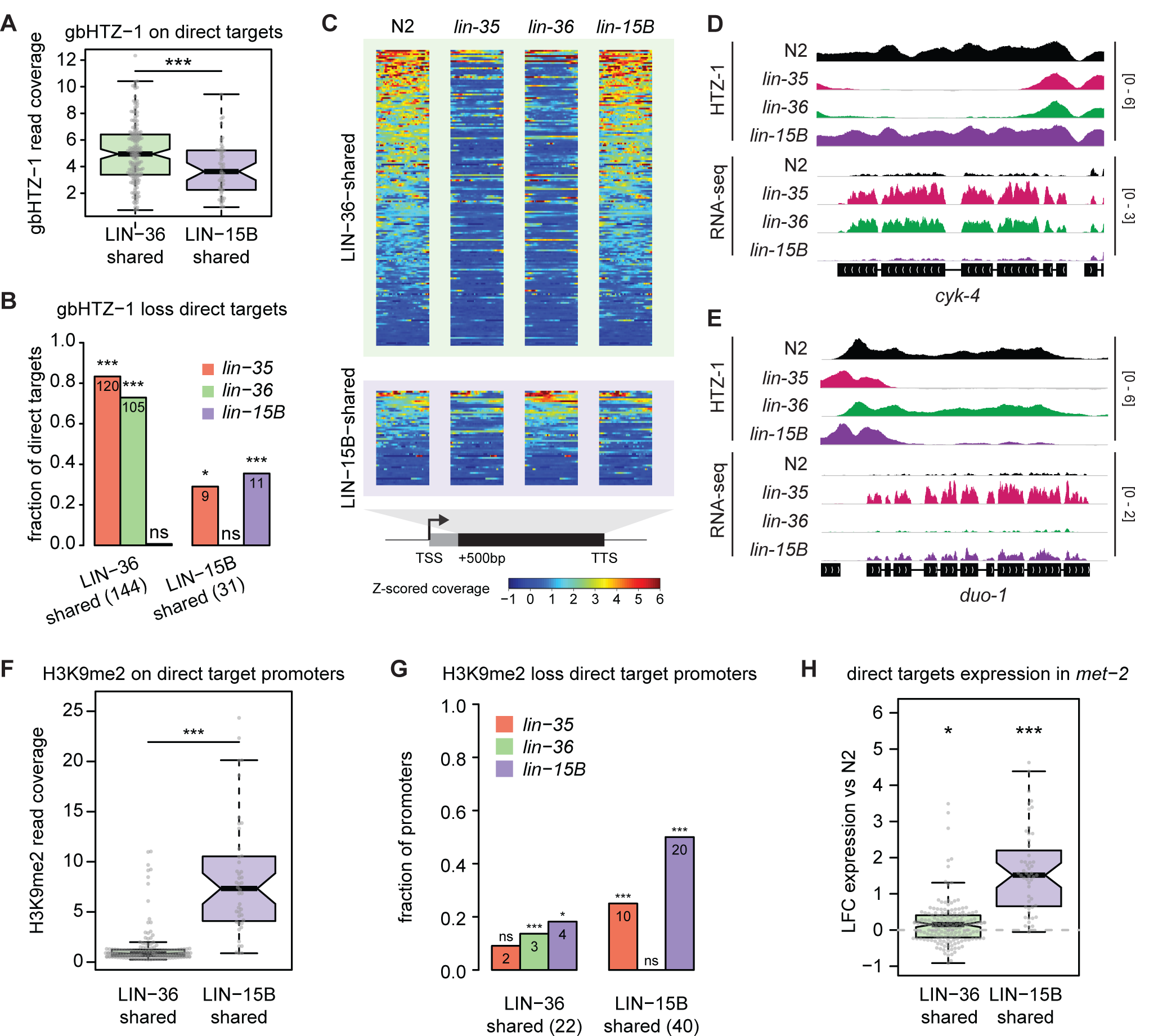
Gene body HTZ-1 and promoter H3K9me2 at LIN-36-shared and LIN-15B-shared targets. **(A)** HTZ-1 coverage over gene bodies of LIN-36-shared and LIN-15B-shared targets. ***P<0.001, Wilcoxon rank sum test. (B) Fraction (and number) of LIN-36-shared and LIN-15B-shared direct targets showing a significant loss of gbHTZ-1 in the respective mutants. *P<0.05, **P<0.01, ***P<0.001, over-representation of gbHTZ-1 loss by hypergeometric test with BH correction. (C) gbHTZ-1 coverage in wild type, lin-35, lin-36 and lin-15B mutants over LIN-36-shared and LIN-15B-shared direct targets. (D) and (E) IGV view of HTZ-1 ChIP-seq and RNA-seq profiles over (D) a LIN-36-shared and (E) a LIN-15B-shared direct target. (F) H3K9me2 coverage at promoters of LIN-36-shared and LIN-15B-shared targets, respectively. ***P<0.001, Wilcoxon rank sum test. (G) Fraction (and number) of LIN-36-shared and LIN-15B-shared direct target promoters showing a significant loss of H3K9me2 in the respective mutants. *P<0.05, ***P<0.001, over-representation of gbHTZ-1 loss by hypergeometric test with BH correction. (H) Log2-fold change of LIN-36-shared and LIN-15B-shared target expression between met-2 mutant and wild-type. ***P< 0.001; *P< 0.05 by t-test.

Evaluating gbHTZ-1 levels on targets in wildtype and mutant starved L1s, we found that the majority of LIN-36-shared targets require both LIN-35 and LIN-36 for high gbHTZ-1 levels, but loss of LIN-15B had no obvious effect at these loci. (Figures 2B-C, S3B; Table S4). In contrast, although some LIN-15B shared targets required LIN-35 and LIN-15B for gbHTZ-1, these were in the minority (Figures 2B, D). Overall, around half (144/293) of all DREAM targets characterised by high gbHTZ-1 correspond to LIN-36-shared targets, and both LIN-36 and LIN-35 function facilitate the recruitment or maintenance of HTZ-1 over these targets.

### LIN-15B promotes H3K9me2 marking for repression of its targets

In addition to differences in gbHTZ-1, we observed a substantial difference in the HTZ-1 profiles over the promoters of different sets of DREAM targets. While LIN-36-shared targets have a bimodal distribution of HTZ-1 flanking the associated LIN-35 and LIN-36 peaks in wild-type animals, HTZ-1 was instead centrally enriched at LIN-15B-shared target peaks (Figure S3C). The HTZ-1 profiles suggest that promoters of LIN-36-shared and LIN-15B-shared targets have different chromatin states. Indeed, whereas LIN-36-shared target peaks showed high DNA accessibility, peaks associated with promoters of LIN-15B-shared targets had low DNA accessibility, indicative of a generally closed chromatin conformation (Figure S3D).

We considered that repression of LIN-15B-shared targets could involve a chromatin-based repression mechanism involving H3K9me2, as previous work showed that LIN-15B facilitates H3K9me2 marking of some DREAM target promoters, although the relevance of H3K9me2 at these genes was not determined (Rechtsteiner et al., 2019). In addition, we observed that LIN-35, LIN-36, and LIN-15B associate with H3K9me2 marked repeats (Figure S1C).

Investigating this connection, we found that H3K9me2 was strongly enriched at LIN-15B-shared, but not LIN-36-shared target promoters (Figure 2F, Table S4). We further found that H3K9me2 marking at LIN-15B-shared target promoters is dependent on LIN-15B (Figure 2G). Notably, H3K9me2 was significantly reduced at 50% of LIN-15B-shared target promoters in lin-15B mutants, and to a lower extent in lin-35 mutants (Figure 2G), whereas little effect was seen in lin-36 mutants or at LIN-36-shared targets.

To test the functional relevance of H3K9me2 in target repression, we profiled gene expression in mutants of met-2, which encodes the major H3K9me2 histone methyltransferase (Bessler et al., 2010). We found that LIN-15B-shared targets had higher expression in met-2 mutants, with 43% being significantly upregulated, whereas met-2 loss had little effect on LIN-36-shared targets (Figure 2H, Table S3). Mechanistically, these results implicate LIN-15B and DREAM in directing repression of their shared targets via MET-2 dependent H3K9me2 promoter marking.

### EFL-1/E2F function is specific for LIN-36-shared targets

We next investigated whether repression of LIN-36-shared and LIN-15B-shared targets differed in their requirement for DREAM components. The DREAM complex consists of DNA binding protein EFL-1/E2F and partner DPL-1/DP1, which are proposed to be bridged to the MuvB sub-complex (LIN-9, LIN-37, LIN-53, LIN-54 and LIN-52) by LIN-35/Rb (Goetsch et al., 2017). To evaluate requirements for different components, we compared gene expression changes among mutants of lin-35/Rb, efl-1, dpl-1, and MuvB sub-complex component lin-37 (Table S3). We found that changes in dpl-1 and lin-37 mutants were similar to those of lin-35 mutants, suggesting a common mechanism. Both LIN-36-shared and LIN-15B-shared targets were derepressed in the two mutants, suggesting that DPL-1 and LIN-37 participate in LIN-35 core roles (Figure S4). In stark contrast, efl-1 mutants only derepressed LIN-36-shared targets (Figure S4). The striking similarities between the lin-36 and efl-1 transcriptomes suggest that EFL-1 functions as a transcriptional repressor specifically at LIN-36-shared DREAM targets.

### E2F motif variants distinguish promoters of LIN-36-shared from LIN-15B-shared targets

To investigate the nature of the differential regulation of the LIN-36-shared versus LIN-15B-shared targets, we searched for DNA sequence motifs that might distinguish their respective promoters (see Methods). We found no motifs specific to either set, however as expected a motif similar to the annotated E2F binding sequence was obtained from searches of each set of direct target promoters (Kirienko and Fay, 2007; Latorre et al., 2015) (Figure S5A). The two E2F motifs showed differences in their consensus sequences, which we named E2F-a (found in LIN-36-shared promoters) and E2F-b (found in LIN-15B-shared promoters). We observed that the strength and enrichment of E2F-a was significantly higher at LIN-36-shared compared to LIN-15B-shared target promoters, and though not significant (Wilcoxon’s rank sum test P = 0.06), there is a trend for E2F-b to be stronger at LIN-15B-shared target promoters (Figures S5B-C). These results suggest that differences in E2F binding sites might explain in part the distinct regulation of LIN-15B-shared and LIN-36-shared target genes.

### LIN-36 and LIN-15B require their THAP domains for function

LIN-36 and LIN-15B both harbor a THAP domain. We assessed the requirements for the THAP domains by creating in-frame deletion alleles (Figure S6A). The LIN-36(ΔTHAP) and LIN-15B(ΔTHAP) mutant proteins were both translated as assessed by western blotting or immunofluorescence (Figures 3B, S6B). We found that deletion of the LIN-36 THAP domain caused loss of nuclear localisation (Figure 3A). In line with this defect, gene expression changes in lin-36(*Δ*THAP) mutants are similar to those of the full deletion mutant (Figure S6C; Table S3). Therefore, the LIN-36 THAP domain is necessary for LIN-36 function, potentially by facilitating nuclear localisation or preventing nuclear export.

**Figure 3.**
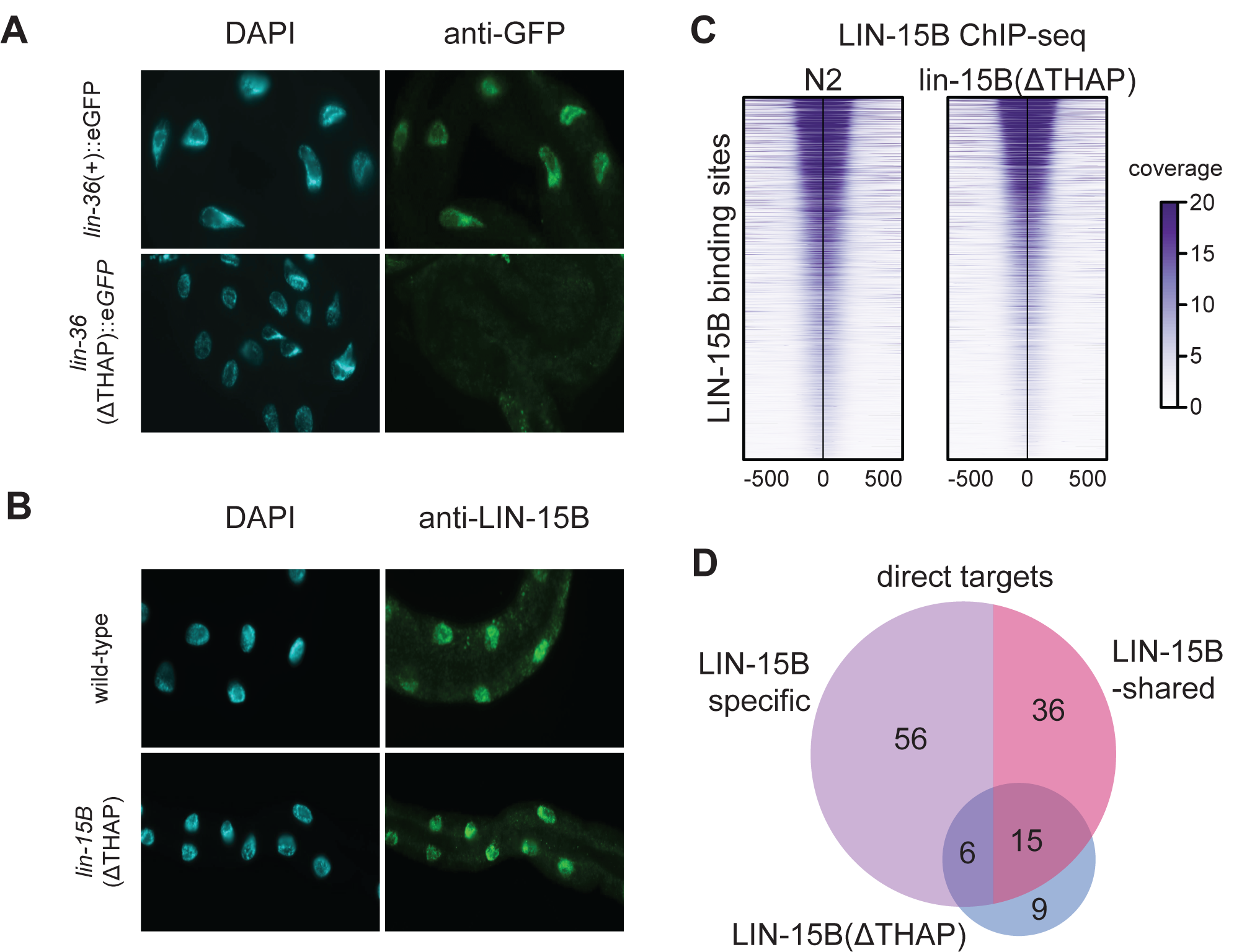
LIN-36 and LIN-15B require their THAP domains for proper function. (A) and (B) Immunofluorescence analysis of LIN-15B and LIN-36 in wildtype and THAP domain deletion mutants. Antibodies used are listed in the Methods section. (C) Heatmap of linear BEADS-normalized LIN-15B ChIP sequencing signal in wild-type and lin-15B(*Δ*THAP) mutant centred over wild-type LIN-15B peaks. (D) Venn diagram of the overlap between direct targets in the indicated mutants. Direct targets shared with LIN-36 were excluded from the total count.

Surprisingly, LIN-15B(ΔTHAP) localized normally to the nucleus and displayed a ChIP binding pattern similar to that of the wild-type protein, with 6774/8861 (∼76%) LIN-15B peaks found in wild-type also called in lin-15B(*Δ*THAP) (Figures 3B-C; Table S2). Despite the relatively normal localisation pattern, 160 genes were derepressed in lin-15B(*Δ*THAP) mutants, including 29% of LIN-15B-shared targets, all of which retained LIN-15B(ΔTHAP) binding (Figures 3D, S6D, Table S3). We conclude that the LIN-15B THAP domain is important but not essential for LIN-15B function. The finding that LIN-15B(ΔTHAP) localises to LIN-15B sites suggests a recruitment mechanism independent of direct DNA binding. LIN-36 and LIN-35 co-facilitate binding, whereas LIN-15B and LIN-35 mutually inhibit binding

### LIN-36 and LIN-35 co-facilitate binding, whereas LIN-15B and LIN-35 mutually inhibit binding

To investigate potential interdependencies in chromatin binding at the LIN-36 and LIN-15B specific targets, we conducted ChIP-seq analyses in mutants (Table S4). We found that LIN-35 and LIN-36 promote the association of EFL-1 and each other at LIN-36-shared targets, with >50% of sites dropping in signal in lin-36 and lin-35 mutants (Figure 4A-B, left panels). In contrast, LIN-36-shared targets showed normal levels of LIN-35, LIN-36, and EFL-1 in lin-15B mutants, consistent with the lack of requirement for LIN-15B at these targets (Figure 5C, left panel). We also found that LIN-15B binding at LIN-36-shared targets was independent of LIN-36 (Figure 4A, left panel). Therefore, LIN-35 and LIN-36 appear to mutually facilitate complex formation and/or stability at LIN-36-shared targets.

**Figure 4.**
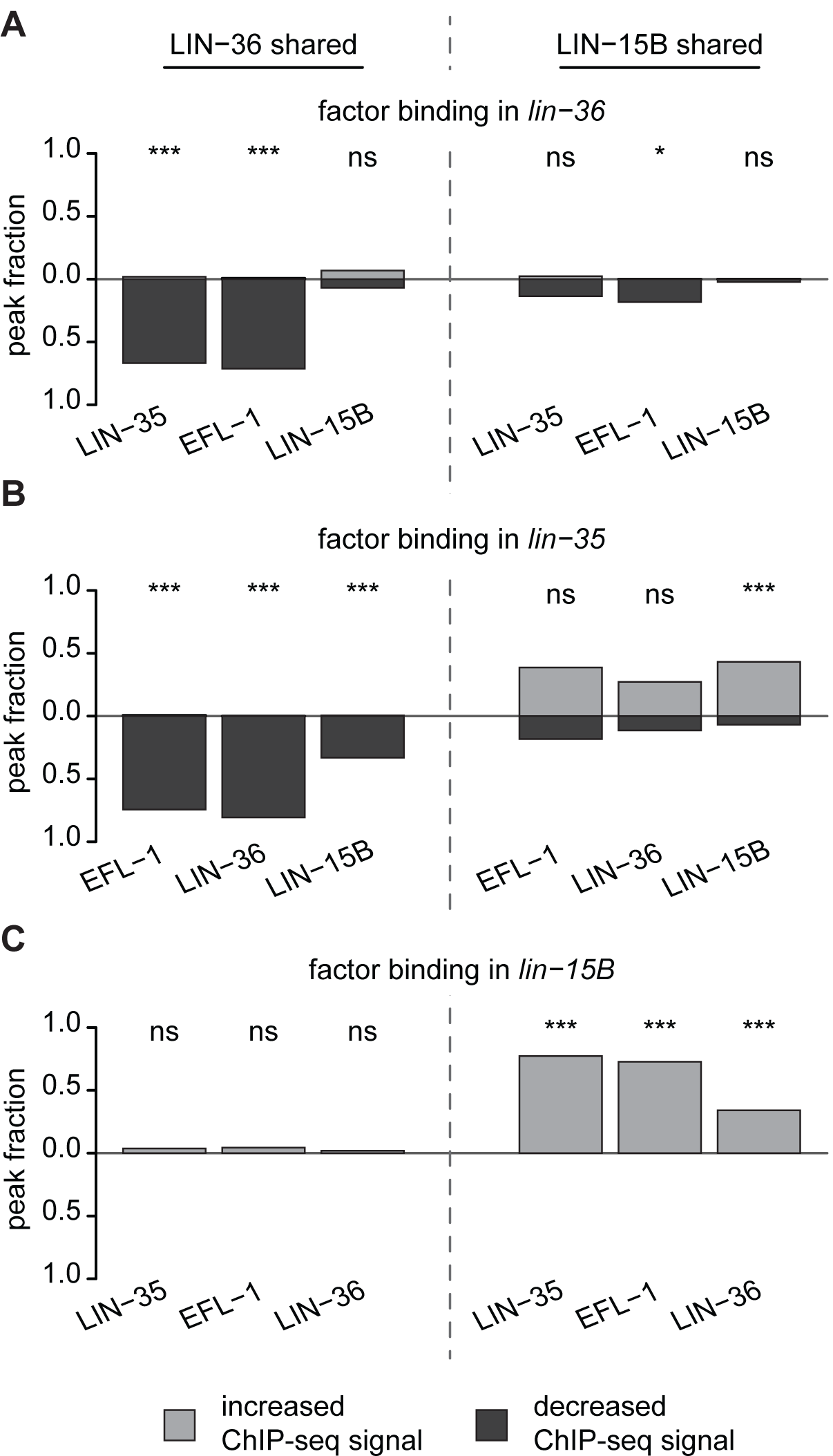
LIN-36 and LIN-35 facilitate, whereas LIN-15B and LIN-35 mutually inhibit each other’s binding. Fraction of LIN-36-shared (left) and LIN-15B-shared (right) promoter-associated peaks showing a significant increase or decrease in ChIP-seq signal in (A) lin-36, (B) lin-35 and (C) lin-15B mutants compared to wt. ***: P<0.001; *: P<0.05; ns: P>0.05, Fisher’s exact test with Benjamini-Hochberg correction.

The LIN-15B-shared targets are strikingly different. At many of these sites, the loss of LIN-15B resulted in an unexpected increase of LIN-35, LIN-36, and EFL-1 signals (Figure 4B, C, right panels). Similarly, lin-35 mutants showed a significant increase in LIN-15B occupancy at LIN-15B shared targets (Figure 5B, C, right panels). In contrast, loss of LIN-36 caused only minor, mostly not significant differences in LIN-35, LIN-15B and EFL-1 binding, (Figure 4A, B, right panel). Intriguingly, we found that the strength of LIN-15B(ΔTHAP) binding was significantly increased at ∼38% of LIN-15B-shared targets (Figure S6E, F), suggesting that the THAP domain may de-stabilise LIN-15B binding. The finding that LIN-35 and LIN-15B repress LIN-15B-shared targets while mutually antagonising chromatin association suggest a potential dynamic cycling of DREAM and LIN-15B which may involve the LIN-15B THAP domain.

## DISCUSSION

The DREAM complex represses cell cycle genes to enforce cellular quiescence, as well as repressing developmental genes to ensure correct patterns of gene expression. While the roles of DREAM have been described in different animals, its mechanism of action is still unclear. Here we show that two THAP domain proteins, LIN-36 and LIN-15B, act with DREAM to repress different sets of target genes through distinct mechanisms.

We found that LIN-36 and LIN-15B bind to thousands of genomic sites shared with LIN-35/Rb. Despite the similarity in binding patterns, genes derepressed upon loss of LIN-36 and LIN-15B are mostly distinct. Consistent with our finding that direct LIN-36 targets are highly enriched for cell-cycle functions, previous work has highlighted a role for LIN-36 in the lin-35 pathway to prevent S-phase entry (Boxem and Van den Heuvel, 2002). Through mutant analyses, we showed that LIN-36 and DREAM mutually stabilise their chromatin association at shared direct targets, and both facilitate high levels of H2A variant HTZ-1 on gene bodies, which we previously found exerts a repressive role on gene expression (Latorre et al., 2015).

The targets that LIN-15B represses with DREAM largely have germline-specific expression. In starved L1 larvae, which are essentially comprised of somatic cells, the promoters of LIN-15B shared-targets have a closed chromatin environment and high levels of H3K9me2. LIN-15B, LIN-35/Rb, and the histone methyltransferase MET-2 are required for H3K9me2 marking and the repression of many of these LIN-15B-shared targets. LIN-35 and LIN-15B ChIP signal at these targets is considerably lower than at LIN-36-shared targets. In contrast to the mutual dependence of LIN-36 and DREAM, LIN-15B and DREAM appear to destabilise each other at shared target promoters. We suggest that mutual destabilisation of LIN-15B and DREAM factors may enable repression by facilitating access of MET-2 and its deposition of repressive H3K9me2.

The presence of a THAP domain in both LIN-36 and LIN-15B suggests a special relationship between DREAM and THAP domain containing proteins. In support of this idea, the human Rb protein shares targets with the THAP1 protein, whose ectopic expression inhibits proliferation in primary human endothelial cells through the transcriptional repression of E2F/Rb targets (Cayrol et al., 2007). Moreover, endogenous THAP1 is necessary for proliferation, suggesting that optimal THAP1 levels are critical. The human THAP11 protein has also been implicated in the regulation of E2F targets and cell proliferation, although its activity is mediated by the interaction with other factors (Brandon Parker et al., 2014). The lack of clear conservation of THAP domain proteins outside this domain suggests that the THAP domain may mediate interactions with DREAM complex. Future work in different systems will further clarify the mechanisms of gene repression employed by the THAP domain protein – DREAM network.

## Supporting information

Code and related files

## ACKNOWLEDGMENTS

The work was supported by an SNF Fellowship to FNC (P400PB_180795) and a Wellcome Trust Senior Research Fellowship to JA (101863). We also acknowledge core support from the Wellcome Trust (092096) and Cancer Research UK (C6946/A14492).

## AUTHOR CONTRIBUTIONS

Conceptualization and methodology: C.G. and J.A.; Software and Formal analysis: C.G., F.N.C. and N.H.; Investigation: C.G., F.N.C., A.A., C.C., Y.D. and J.M.; Writing – original draft preparation: C.G., F.N.C. and J.A.; Supervision and Funding acquisition: J.A.

## DECLARATION OF INTERESTS

The authors declare no competing interests.

## Methods

### Worm culture and strains

Strains were cultured using standard methods (Brenner, 1974). Strains used in the paper are given in Table S1.

### Generation of psep-1::his-58::eGFP, lin-36::eGFP, lin-36 deletion, and THAP domain deletion alleles

The psep-1::his-58::eGFP transgene was generated using three-site Gateway cloning (Invitrogen) in the MosSCI compatible vector pCFJ150, which targets Mos site Mos1(ttTi5605) (Frøkjær-Jensen et al., 2008). The sep-1 promoter (chr I: 3439109-3438531) was cloned into site one. Plasmids pJA273 and pJA257 (Zeiser et al., 2011) were used to put his-58 into site 2 and eGFP::tbb-2-3’UTR into site three, respectively. MosSCI lines were generated as described (Frøkjær-Jensen et al., 2008).

CRISPR-Cas9 genome editing was used to generate the following strains: JA1798: lin-15B(we23[*Δ*THAP]) X, JA1810: lin-36(we30[lin-36::eGFP]) III, JA1811: lin-36(we30[lin-36::eGFP], we31[*Δ*THAP]) III, and JA1850: lin-36(we36) III (Table S1). Injections were performed using gRNA-Cas9 ribonucleoprotein (RNP) complexes preassembled in vitro (Paix et al., 2017). dpy-10 co-CRISPR method was used to enrich for desired edit (Arribere et al., 2014; Paix et al., 2015). Cas9 protein was made in-house (Paix et al., 2015); tracrRNA and crRNAs were purchased from Dharmacon or Integrated DNA Technologies; repair templates were purchased from IDT as Ultramer oligonucleotides; eGFP double stranded amplicons were generated by standard PCR (Paix et al., 2017). crRNAs were designed using the online CRISPOR tool (Haeussler et al., 2016). JA1798, JA1810 and JA1850 were made in the Bristol wild-type N2 background; JA1811 was made in JA1810.

### RNAi Screen

An RNAi sub-library targeting 1104 known or predicted nuclear proteins was used for the RNAi (Table S1). The primary screen was carried out in four replicates, two feeding from the L3 stage and two feeding from the YA stage, the latter to avoid the high embryonic lethality induced by some clones. Bacteria were grown at 37°C overnight in 900 μl LB (supplemented with 10 μg/ml carbenicillin, 10 μg/ml tetracyline, and 100 U/ml nystatin) in 96 well plates. RNAi expression was induced through the addition of 4 mM IPTG, and bacteria were further incubated for 3 hours at 37°C. Bacteria were then pelleted and resuspended in 450 μl of S medium (Stiernagle, 2006), 50 μl was transferred into each well of a new 96-well plate, and approximately 10-15 L3 or YA psep-1::his-58::eGFP animals were placed into each well. The animals were monitored and when most had L1 progeny the L1s were analysed for increased expression of the reporter using a COPAS (Union Biometrica) profiler by measuring fluorescence intensity of L1 sized progeny. In the primary screen, 210 clones induced de-repression of the reporter in two out of the four replicates and were included in four replicates of a secondary screen conducted using YA aninals. Of these, 36 showed de-repression in three out of four replicates and were considered to be hits (see Table S1). These clones were sequenced and verified.

### RNAi screen of THAP genes

RNAi plates targeting THAP domain genes were prepared as in (Ahringer, 2006). Synchronized L3 psep-1::his-58::eGFP animals were placed onto RNAi plates and their progeny assessed daily for somatic GFP expression through visual observation under a fluorescent microscope, qualitatively compared to control RNAi. Experiments were carried out three times.

### Collection of starved L1 animals for RNA-seq and ChIP-seq

Synchronized adults were grown at 20°C in liquid culture using standard S-basal medium and HB101 E. coli, bleached to isolate embryos, the eggs hatched 20-22 hours at 25°C in M9 buffer, and then the starved L1s were sucrose floated and collected by flash freezing in liquid nitrogen. The efl-1(se1ts) mutants were hatched at 26°C (Page et al., 2001).

### ChIP-seq

Frozen starved L1 worms were ground to a powder, which was incubated in 1.5 mM EGS (Pierce 21565) in PBS for 8 minutes, followed by the addition of formaldehyde to a final concentration of 1%, and incubated for a further 8 minutes. The fixation was quenched for 5 minutes by the addition of 0.125 M glycine. Fixed tissue was washed 2X with PBS with protease inhibitors (Roche EDTA-free protease inhibitor cocktail tablets 05056489001) and once in FA buffer (50 mM Hepes pH7.5, 1 mM EDTA, 1% TritonX-100, 0.1% sodium deoxycholate, and 150 mM NaCl) with protease inhibitors (FA+), then resuspended in 1 ml FA+ buffer per 1 ml of ground worm powder. The extract was sonicated to an average size of ∼250 base pairs using a Bioruptor Pico (Diagenode), and 10-20 ug of DNA was used per ChIP reaction, together with ∼1ug DNA from C. briggsae ChIP extract. Antibodies used for ChIP are provided in Table S1 and ChIP-seq datasets are described in Table S5. ChIP and library preparations were done as described in (Jänes et al., 2018).

### RNA-seq

A single ball of frozen worms was used for RNA extractions. Total RNA was extracted using TriPure (Roche) and further purified using an RNeasy column (Qiagen). RNA-seq libraries were prepared from 100-1000 ng of total RNA using the Illumina TruSeq RNA kit according to the manufacturers’ instructions. RNA-seq datasets are given in Table S5.

### Data processing

ChIP-seq and RNA-seq libraries were sequenced using Illumina HiSeq1500. ChIP-seq reads were aligned to a concatenated WS235/ce11 + cb3 assembly of the C. elegans and C. briggsae genomes using BWA v. 0.7.7 with default settings (BWA-backtrack algorithm) (Li and Durbin, 2010), but only C. elegans data were analysed here. The SAMtools v. 0.1.19 ‘view’ utility (Li et al., 2009) was used to convert the alignments to BAM format. Normalized mapq10 ChIP-seq coverage tracks were generated using the BEADS algorithm (Cheung et al., 2011). RNA-seq reads were aligned using STAR (Dobin et al., 2013) with the two-pass mode using the C. elegans gene annotation from Wormbase (version WS260) as a guide (after removing any gene annotation from the mitochondrial DNA). BigWig tracks were generated using the wigToBigWig tool downloaded from the UCSC website (http://hgdownload.soe.ucsc.edu/).

### Differential expression analysis

A gene model was built based on the WS260 annotation. Tag counts for each gene were extracted from STAR aligned BAM files, and differential gene expression between N2 and mutant backgrounds was tested using DESeq2 (Love et al., 2014). A false discovery rate (FDR) < 0.01 and LFC > 0.5849 was used to define genes as up-regulated, and FDR < 0.01 and LFC < −1 was used to define genes as down-regulated. Table S3 contains the DESeq2 log2 fold change and FDR for each mutant vs. wildtype comparison.

### Peak calls and annotation to genes

ChIP-seq peaks were called for each factor using YAPC (https://github.com/jurgjn/yapc) (Jänes et al., 2018). Briefly, peak calls were generated through identification of concave regions (regions with negative smoothed second derivative) using the BEADS normalized bigwig tracks. The candidate peaks were tested for statistical significance between replicates using IDR (Li et al., 2011), and only peaks with FDR <0.001 were kept in our datasets. For three factors (LIN-35, LIN-15B, and EFL-1) we had validated antibodies against the protein; however, to determine LIN-36 binding, we endogenously CRISPR tagged it using GFP. To test that the GFP tag did not disturb the binding of the other factors, we also chromatin immunoprecipitated the other factors in the lin-36::eGFP strain. For each factor the Spearman correlation (calculated using DeepTools (Ramírez et al., 2016)) over the peak calls is between 0.76 and 0.98 (Table S2). Therefore, to call wildtype peaks for LIN-35, LIN-15B and EFL-1 we have used all four of our biological replicates, whilst we have used two for LIN-36. We further redefined these calls by merging overlapping LIN-35, LIN-36 and/or LIN-15B peaks, and then re-scaling merged and factor specific peaks to +/-100bp around their midpoint. The resulting peaks were assigned to genes, if they were within the furthest upstream promoter (Jänes et al., 2018) and the end of the gene (Table S2). Peak overlap with regulatory elements or Dfam2.0 annotated repeats (Hubley et al., 2016) was determined using BEDTools (Quinlan and Hall, 2010).

### Identification of direct targets

Direct targets of a given protein were defined as genes up-regulated in a mutant condition and that have an associated peak for that protein. The LIN-36-shared and LIN-15B-shared direct targets are direct targets of both LIN-36 and LIN-35, or both LIN-15B and LIN-35, respectively, but not upregulated in lin-15B or lin-36, respectively (Table S3).

### GO enrichment analysis

Enrichment for specific gene ontology terms was obtained using the Gene Enrichment Analysis (GEA) tool (Angeles-Albores et al., 2018) available on Wormbase.

### Gene body HTZ-1 enrichment

Average gene body HTZ-1 (gbHTZ-1) read coverage was calculated from the region from the most upstream Wormbase TSS +500bp to the most downstream TTS. We identified genes showing a significant loss of HTZ-1 (LFC vs N2 < 0, adjusted P < 0.001) by running DESeq2 on the coding genes in the top 90% of gbHTZ-1 coverage in N2. Genes shorter than 500 bp in length were excluded from the analysis.

### H3K9me2 enrichment

Average H3K9me2 signal (BEADS normalized linear coverage) was calculated over LIN-35 + LIN-36 or LIN-35 + LIN-15B ChIP peaks associated to the putative promoter region (i.e. −500 – 0bp upstream of any Wormbase coding transcript) of their respective LIN-36-shared or LIN15B-shared direct targets. Peaks showing a significant loss of H3K9me2 (LFC vs N2 < 0, adjusted P < 0.01) were identified using DESeq2 on LIN-35, LIN-36 and/or LIN-15B peaks overlapping a wild-type H3K9me2 peak (called using MACS (Zhang et al., 2008) with standard settings).

### Motif enrichment analysis

DNA motifs enriched at LIN-36-shared and LIN-15B-shared promoter-associated peaks were detected using MEME (Bailey et al., 2009) (with: -objfun de). Enriched motifs were then re-annotated using FIMO (with: --thresh 1e-2). Similarity to known motifs was evaluated using TOMTOM.

### Identification of differentially bound peaks

DESeq2 was used to identify peaks differentially bound between wild type and a mutant background by comparing the read counts from the bwa aligned BAM files mapped in wild-type peak regions. Peaks with increased signal in mutants have adjusted P-value < 0.001 and LFC > 0. Peaks with decreased signal in mutants have adjusted P < 0.001 and LFC < 0.

### Data and Code availability

Raw and processed data generated during this study are available at NCBI Gene Expression Omnibus (GEO) under accession code GSE155191. Processing of genomic coordinates was performed using the BEDTools suite (version 2.27.1) and in-house scripts. Statistical analyses were performed in R (Yan et al., 2011). Commands used to process data are available as Supplementary file.

## SUPPLEMENTARY FIGURE LEGENDS

**Figure S1. LIN-35, LIN-36 and LIN-15B co-localize extensively on chromatin.**

(A) Heatmaps of BEADS normalized linear ChIP tracks centred over the indicated regions. Tracks are of combined replicates. We note that signal at single factor sites is generally weak and therefore confidence that other factors are not present is not high. (B) Assignments of peaks to features in the genome. Peaks were first overlapped with regulatory elements identified in Janes et al. 2018, then with repetitive elements from Dfam2.0. (C) Fraction of LIN-35, LIN-15B and LIN-36-bound repeats (from Dfam 2.0) showing enrichment for H3K9me2.

**Figure S2. Expression and binding profile of LIN-35, LIN-36 and LIN-15B targets.**

(A) Expression level (measured as log2 TPM) of genes upregulated in lin-35, lin-36(we36), and lin-15B in the germline and in different types of dividing (SGP: somatic gonad precursors, SC: seam cells, I: intestine) and non-dividing (NSH: non-seam hypodermis, BWM: body wall muscle) cell types. The dashed grey line indicates a TPM value of 1. Expression difference between germline and any other tissue was significant for LIN-35, LIN-36 and LIN-15B targets (Benjamini-Hochberg adjusted Mann-Whitney test P<10^−3^) (B) Germline expression specificity (calculated as expression in germline / sum of expression in any cell type) of LIN-35-specific, LIN-36-shared and LIN-15B-shared direct targets. (C) and (D) Heatmaps of BEADS normalized linear ChIP tracks centred over the LIN-35+LIN-36 and LIN35+LIN-15B peaks associated to the promoters of LIN-36-shared (C) and LIN-15B-shared (C and D, in different scales) direct targets. Significant differences (Wilcoxon rank sum test): ***: P<0.001. Data for panels (A, B) are from Cao et al, 2017.

**Figure S3. Different silencing mechanisms of LIN-36-shared targets and LIN-15B-shared targets.**

(A) fraction of LIN-35-specific, LIN-36-shared and LIN-15B-shared direct targets with high (dark grey) or low (light grey) levels of gbHTZ-1. (B) Number of coding genes showing a significant reduction in gbHTZ-1 levels in the respective mutants. Dark grey bars indicate direct targets. (C) Signal plot of Z-scored HTZ-1 coverage calculated over the LIN-35+LIN-36 and LIN35+LIN-15B peaks associated to the promoters of LIN-36-shared (green) and LIN-15B-shared (purple) direct targets. (D) Signal plot of ATAC-seq signal (in RPM coverage) from L1-staged larvae over the LIN-35+LIN-36 and LIN35+LIN-15B peaks associated to the promoters of LIN-36-shared (green) and LIN-15B-shared (purple) direct targets. Data from Janes et al., 2018. (E) Number of LIN-35, LIN-36 and/or LIN-15B peaks showing a significant reduction in H3K9me2 levels in mutants.

**Figure S4. Distinct set of direct targets are upregulated in other DREAM factor mutants.**

Fraction of LIN-36-shared (green) and LIN-15B-shared (purple) direct targets showing upregulated expression in dpl-1, efl-1 and lin-37 mutants. Significant differences (LIN-36-shared vs LIN-15B-shared fraction, Fisher’s exact test with Benjamini-Hochberg correction): ***: P<0.001; ns: P>0.05.

**Figure S5. E2F binding motif variants enriched at LIN-36-shared and LIN-15B-shared target promoters.**

(A) Motif E2F-a was derived from LIN-36-shared sites and E2F-b from LIN-15B shared sites. (B) Fraction of LIN-36-shared and LIN-15B-shared promoter-associated peaks bearing a strong (FIMO P < 0.001) E2F-a or E2F-b variant. Significant differences (Fisher’s exact test with Benjamini-Hochberg correction): *: P < 0.05; ns: P > 0.05. (C) strength of E2F-a and E2F-b motifs found at LIN-36-shared and LIN-15B-shared promoter-associated peaks. Significant differences (Wilcoxon rank sum test): ***: P < 0.001; ns: P > 0.05.

**Figure S6. Effects of THAP domain deletion in LIN-36 and LIN-15B.**

(A) Diagram of the LIN-15B and LIN-36 proteins, illustrating their THAP domains and the deletions generated in this study. Arrows indicate the positions of the premature stop codons in the corresponding alleles. (B) Western blot of LIN-36 and LIN-36(ΔTHAP). Anti-GFP antibody was used to detect both proteins. (C) and (D) Overlap between genes upregulated in (C) lin-36 or (D) lin-15B mutant strains used in this study. (E) IGV view of a representative LIN-15B-shared direct target. Factor-specific ChIP-seq enrichment shown as linear BEADS-normalized tracks. RNA sequencing data depict read-depth normalized combined replicates. (F) Fraction of LIN-36-shared (left) and LIN-15B-shared (right) promoter-associated LIN15B(ΔTHAP) peaks showing a significant difference in occupancy in lin-15B(ΔTHAP) mutants. Significant differences (up-vs downregulated fraction, Fisher’s exact test with Benjamini-Hochberg correction): ***: P < 0.001; ns: P > 0.05.

## Notes

### Competing Interest Statement

The authors have declared no competing interest.

