## Supplementary figures and images for "DREAM represses distinct targets by cooperating with different THAP domain proteins"

### Supplementary Figure S1

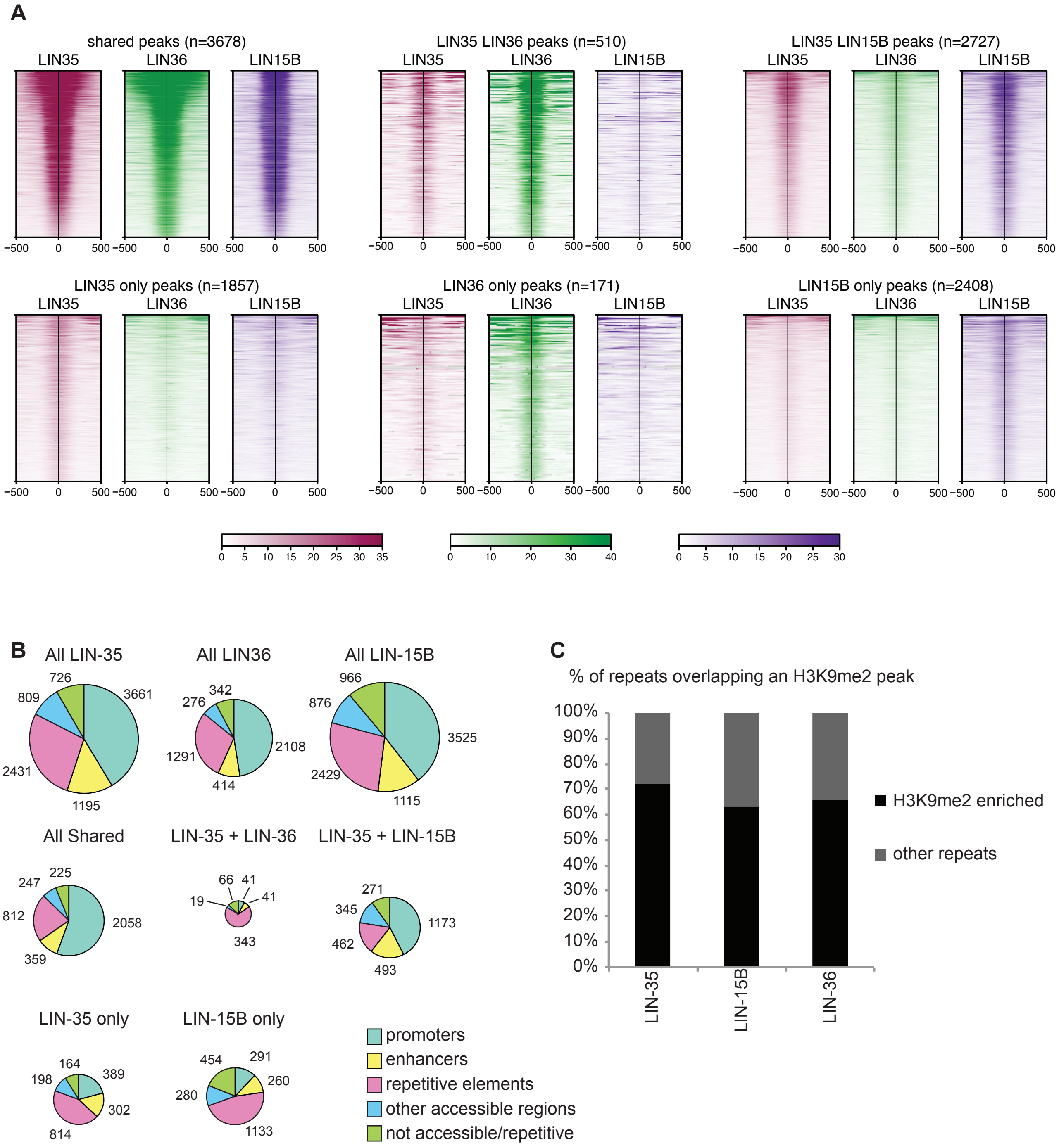

### Supplementary Figure S2

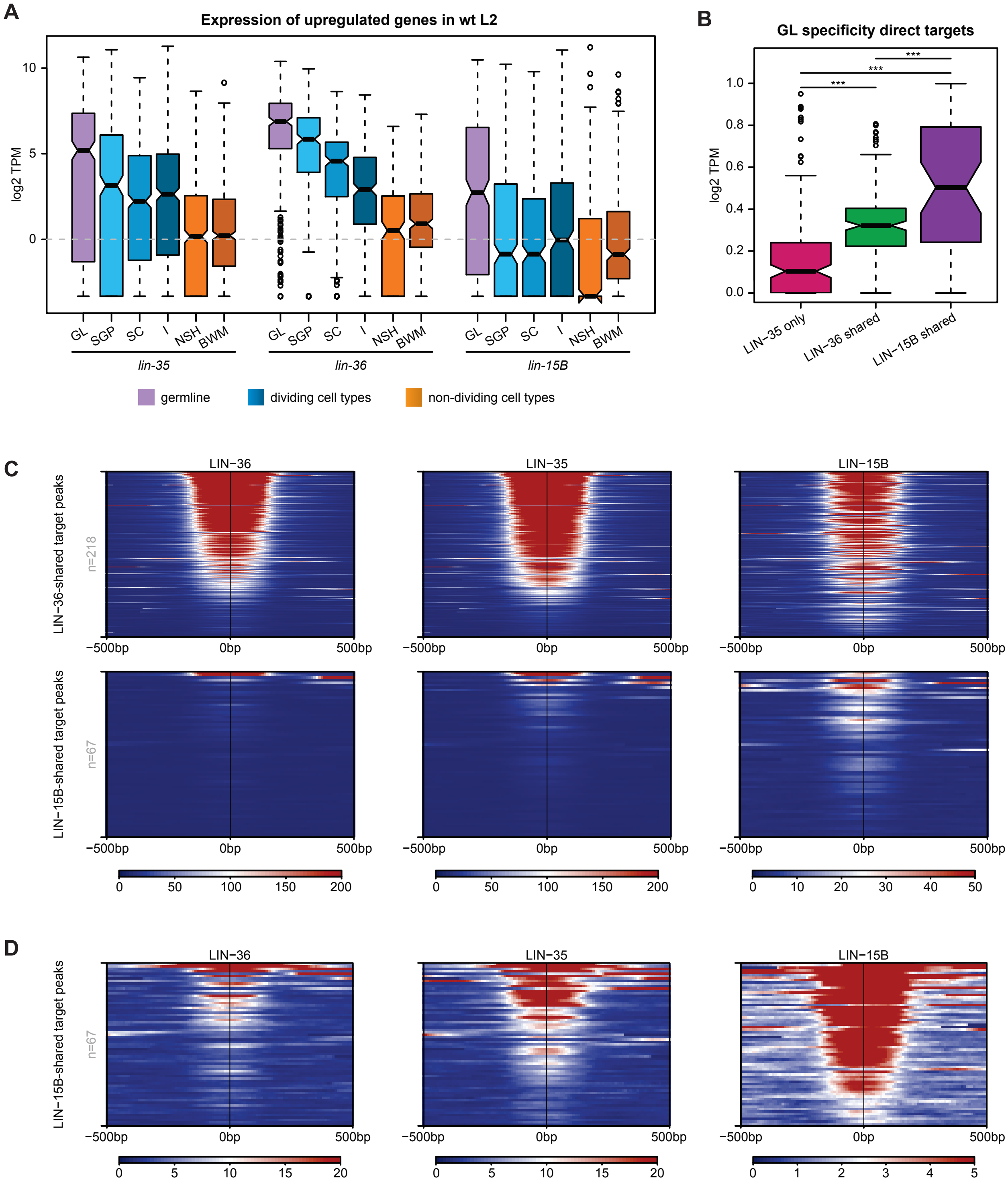

### Supplementary Figure S3

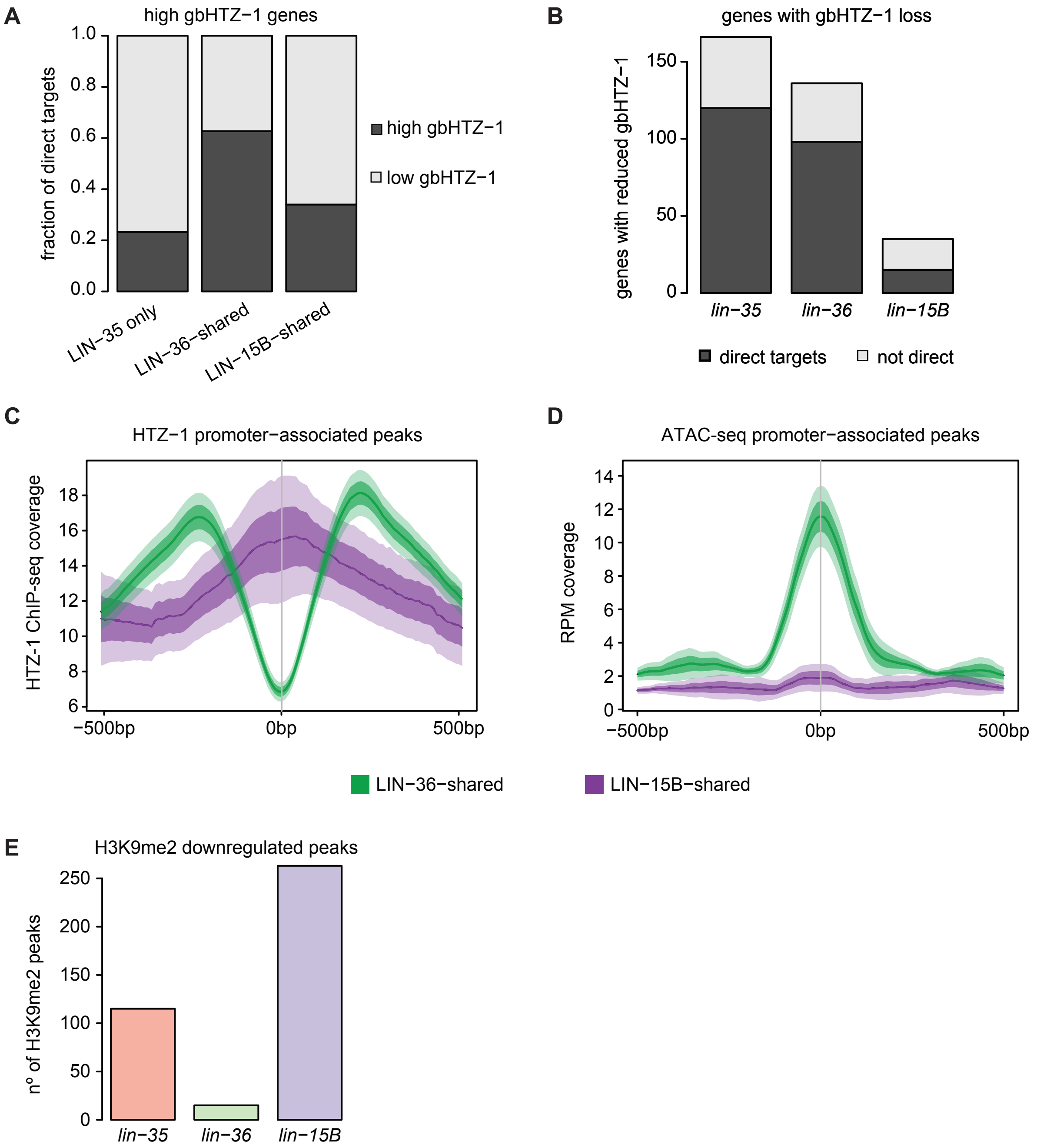

### Supplementary Figure S4

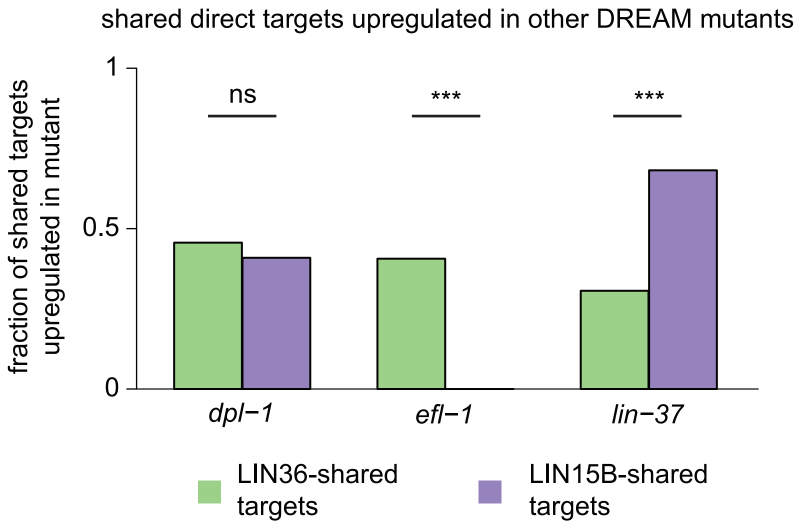

### Supplementary Figure S5

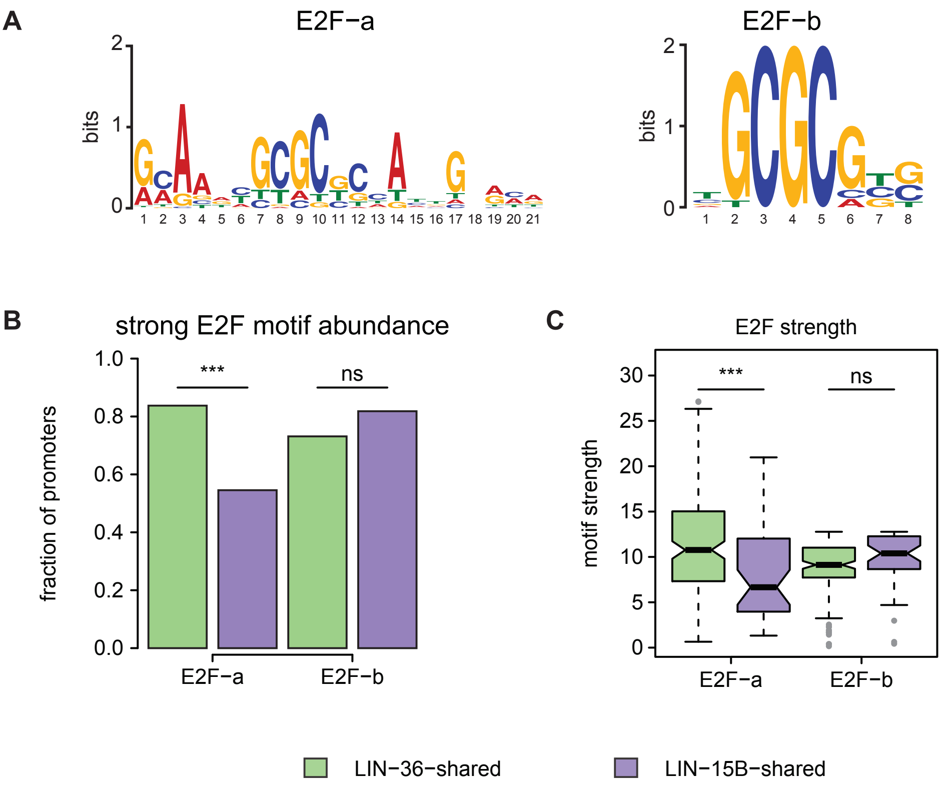

### Supplementary Figure S6

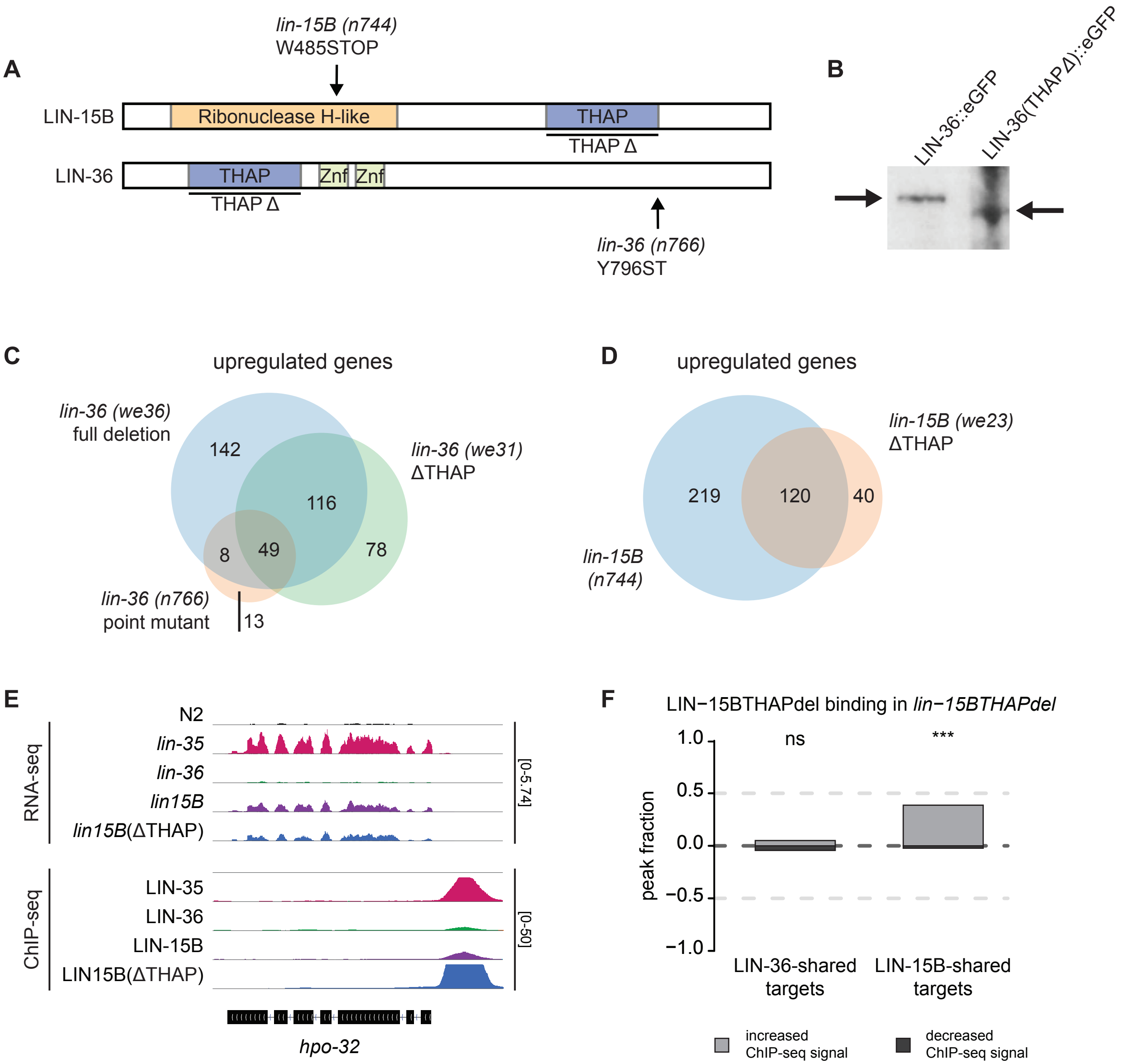
