## Supplementary material for "DREAM represses distinct targets by cooperating with different THAP domain proteins": Code and related files: bwa_alignment.sh

```
#!/bin/bash
#
#SBATCH --job-name=bwa
#SBATCH --output=/mnt/home1/ahringer/fnc21/slurm_out/bwa.%N.%j.out
#SBATCH --error=/mnt/home1/ahringer/fnc21/slurm_err/bwa.%N.%j.err
#SBATCH --ntasks=1
#SBATCH -N 1
#SBATCH -n 8
#SBATCH --mem-per-cpu=4000
#SBATCH --mail-type=END,FAIL
#SBATCH --mail-user=

working_dir=$1
input_filename=$2
chip=$3
species=$4
macs=$5


genome_idx=$working_dir/bwa_idx/$species
output_path_name=$working_dir/data/$species/$chip/alignment
genome_path=$working_dir/species/$species/genome
fastq_path=$working_dir/data/$species/$chip/fastq
macs_output_dir=$working_dir/data/$species/$chip/peaks/$input_filename

mkdir -p $output_path_name

bwa aln -t 10 $genome_idx $fastq_path/$input_filename.fq.gz > $output_path_name/$input_filename.sai
bwa samse -f $output_path_name/$input_filename.sam $genome_idx $output_path_name/$input_filename.sai $fastq_path/$input_filename.fq.gz

samtools view -@ 8 -bT $genome_path/$species.fa $output_path_name/$input_filename.sam > $output_path_name/$input_filename.bam
rm $output_path_name/$input_filename.sam
rm $output_path_name/$input_filename.sai

samtools sort -@ 8 $output_path_name/$input_filename.bam > $output_path_name/$input_filename.sorted.bam
rm $output_path_name"/"$input_filename.bam
samtools index $output_path_name/$input_filename.sorted.bam

samtools view -@ 8 -b -q 10 $output_path_name/$input_filename.sorted.bam > $output_path_name/$input_filename.sorted.mapq10.bam
samtools index $output_path_name/$input_filename.sorted.mapq10.bam

if [ $macs == "yes" ]
then
    mkdir $macs_output_dir
    macs2 callpeak --bdg --SPMR --gsize ce --nolambda -n $input_filename --outdir $macs_output_dir -t $output_path_name/$input_filename.sorted.bam
    sort -k 1,1 -k2,2n $macs_output_dir"/"$input_filename"_treat_pileup.bdg" > $macs_output_dir"/"$input_filename"_treat_pileup.sorted.bdg"
    bedGraphToBigWig $macs_output_dir"/"$input_filename"_treat_pileup.sorted.bdg" $genome_path/$species.chrom.sizes.txt $macs_output_dir"/"$input_filename"_treat_pileup.sorted.bw"
fi
```
