## Supplementary material for "DREAM represses distinct targets by cooperating with different THAP domain proteins": Code and related files: DESeq_all_mutants.R

```
library("DESeq2")
library("GenomicAlignments")
library("rtracklayer")

args <- commandArgs()
main_working_dir = args[6]

working_dir=paste(main_working_dir, "/DR_peaks/", sep="")

#Import the gene model and the bam files that I want the counts to be done on 
sample_ID_list=read.table(paste(main_working_dir, "/chip_seq_samples.txt", sep=""), sep=".", col.names=c("chip", "strain", "replicate", "sample"))
TFs=levels(sample_ID_list$chip)

for (TF in TFs){
  dir.create(file.path(main_working_dir, "DR_peaks", TF))
  TF_ID_list = sample_ID_list[sample_ID_list$chip == TF,]
  TF_ID_list$strain <- factor(TF_ID_list$strain)
  bam_files = c()
  bam_files_samples=c()
  for (i in c(1:dim(TF_ID_list)[1])){
    sampleID=TF_ID_list[i,4]
    strain=TF_ID_list[i,2]
    bam_files_sample_name=paste(strain, sampleID, sep=".")
    bam_file_sample=grep(sampleID, list.files(path=paste(main_working_dir, "/data/elegans/chipseq/alignment", sep=""), pattern = "\\.sorted\\.bam$", full.names = T), value=T)
    bam_files = c(bam_files, bam_file_sample)
    bam_files_samples = c(bam_files_samples, bam_files_sample_name)
  }
  peaks <- import.bed(paste(main_working_dir, "/peaks/wt_rescaled/", TF, "_all.peaks.bed", sep=""))
  peaks_tab <- read.table(paste(main_working_dir, "/peaks/wt_rescaled/", TF, "_all.peaks.bed", sep=""), sep = "\t", col.names = c("chrom", "start", "end", "peak_id"))
  bf = BamFileList(bam_files)
  se <- summarizeOverlaps(features=peaks, bf, mode = "Union")
  
  colData(se)$strain <-  factor(TF_ID_list$strain)
  de <- DESeqDataSet(se, ~strain)
  de1 <- DESeq(de)
  
  de_est <- estimateSizeFactors(de)
  normalized_counts <- counts(de_est, normalized=TRUE)
  colnames(normalized_counts) = bam_files_samples
  normalized_counts_complete=cbind(peaks_tab, normalized_counts)
  write.table(normalized_counts_complete, file=paste(working_dir, TF, "/" , TF, "_normalized_counts_DESeq.txt", sep=""), sep="\t", quote=F, row.names=F)
  
  all_strains=levels(TF_ID_list$strain)
  all_mutants=all_strains[all_strains != "N2" & all_strains != "lin36::eGFP"]
  for (mutant in all_mutants){
    # results against N2 wt
    if ("N2" %in% all_strains){
      res <- results(de1, c('strain', mutant, 'N2'))
      res_peak=as.data.frame(cbind(peaks_tab, res))
      downreg=res_peak[res_peak$log2FoldChange < 0 & res_peak$padj < 0.001 & !is.na(res_peak$padj),]
      upreg=res_peak[res_peak$log2FoldChange > 0 & res_peak$padj < 0.001 & !is.na(res_peak$padj),]
      dir.create(file.path(working_dir, TF))
      write.table(downreg, file = paste(working_dir, TF, "/" , TF, "_downregulated_", mutant, "_vs_N2.txt", sep=""), quote=F, row.names = F, sep="\t")
      write.table(upreg, file = paste(working_dir, TF, "/" , TF, "_upregulated_", mutant, "_vs_N2.txt", sep=""), quote=F, row.names = F, sep="\t")
      write.table(res_peak, file = paste(working_dir, TF, "/" , TF, "_DESeq_", mutant, "_vs_N2.txt", sep=""), quote=F, row.names = F, sep="\t")}
    # results against lin36GFP wt
    if ("lin36::eGFP" %in% all_strains){
      res <- results(de1, c('strain', mutant, 'lin36::eGFP'))
      res_peak=as.data.frame(cbind(peaks_tab, res))
      downreg=res_peak[res_peak$log2FoldChange < 0 & res_peak$padj < 0.001 & !is.na(res_peak$padj),]
      upreg=res_peak[res_peak$log2FoldChange > 0 & res_peak$padj < 0.001 & !is.na(res_peak$padj),]
      dir.create(file.path(working_dir, TF))
      write.table(downreg, file = paste(working_dir, TF, "/" , TF, "_downregulated_", mutant, "_vs_lin36::eGFP.txt", sep=""), quote=F, row.names = F, sep="\t")
      write.table(upreg, file = paste(working_dir, TF, "/" , TF, "_upregulated_", mutant, "_vs_lin36::eGFP.txt", sep=""), quote=F, row.names = F, sep="\t")
      write.table(res_peak, file = paste(working_dir, TF, "/" , TF, "_DESeq_", mutant, "_vs_lin36::eGFP.txt", sep=""), quote=F, row.names = F, sep="\t")}
  }
}
```
