## Supplementary material for "DREAM represses distinct targets by cooperating with different THAP domain proteins": Code and related files: DESeq_gene_exp.R

```
library("DESeq2")
library("GenomicAlignments")
library("rtracklayer")
library("GenomicFeatures")

args <- commandArgs()

main_working_dir = args[6]
dir.create(file.path(main_working_dir, "DE_genes"))

working_dir=paste(main_working_dir, "DE_genes/", sep="/")

#Import the gene model and the bam files that I want the counts to be done on 
sample_ID_list=read.table(paste(main_working_dir, "RNASeq_samples.txt", sep="/"), col.names = c("sample", "strain"))
strains=levels(sample_ID_list$strain)

bam_files=c()
for (i in c(1:dim(sample_ID_list)[1])){
    bam_files = c(bam_files, paste(main_working_dir, "/data/elegans/RNASeq/alignment/", sample_ID_list[i,2], "/", sample_ID_list[i,2], ".", sample_ID_list[i,1], ".Aligned.sortedByCoord.out.bam", sep=""))
}

bf = BamFileList(bam_files )
gtf = import(paste(main_working_dir, "/species/elegans/gene_annotation/elegans.annotations.coding_genes.ce11.gtf", sep=""))
gtf = reduce(split(gtf, gtf$gene_name))
genehits <- summarizeOverlaps(gtf, bf, mode="Union")
colData(genehits)$strain = factor(sample_ID_list$strain)
de <- DESeqDataSet(genehits, ~strain)
de1 <- DESeq(de)


de_est <- estimateSizeFactors(de)
normalized_counts <- counts(de_est, normalized=TRUE)
write.table(normalized_counts, file=paste(working_dir, "normalized_counts_DESeq.txt", sep=""), sep="\t", quote=F, col.names=NA)

current_date=gsub("-", "", scan(text = as.character(Sys.time()), what = "")[1])
current_time=gsub(":", "", scan(text = as.character(Sys.time()), what = "")[2])

all_strains=levels(sample_ID_list$strain)
all_mutants=all_strains[all_strains != "N2" & all_strains != "N2_26C"]
for (mutant in all_mutants){
    if (mutant == "efl1") {
        res <- results(de1, c('strain', mutant, 'N2_26C'))
    }
    else {
        res <- results(de1, c('strain', mutant, 'N2'))
    }
    downreg=res[res$log2FoldChange <= -1 & res$padj < 0.01 & !is.na(res$padj),]
    upreg=res[res$log2FoldChange >= 0.5849 & res$padj < 0.01 & !is.na(res$padj),]
    #pdf(paste(main_working_dir, "/plots/MA_plots/MA_plot_", mutant, ".gene_exp.", current_date, ".", current_time,".pdf", sep=""), width=8, height=8)
    #plotMA(res, ylim=c(-2,2))
    #dev.off()
    write.table(downreg, file = paste(working_dir, "genes_downregulated_", mutant, "_vs_N2.txt", sep=""), quote=F, col.names = F, row.names = T, sep="\t")
    write.table(upreg, file = paste(working_dir, "genes_upregulated_", mutant, "_vs_N2.txt", sep=""), quote=F, col.names = F, row.names = T, sep="\t")
    write.table(res, file = paste(working_dir, "genes_DESeq_", mutant, "_vs_N2.txt", sep=""), quote=F, row.names = T, sep="\t")
}
```
