## Supplementary material for "DREAM represses distinct targets by cooperating with different THAP domain proteins": Code and related files: DESeq_HTZ1.R

```
library("DESeq2")
library("GenomicAlignments")
library("rtracklayer")

args <- commandArgs()

main_working_dir = args[6]

working_dir=paste(main_working_dir, "HTZ1/", sep="/")

#Import the gene model and the bam files that I want the counts to be done on 
TF_ID_list=read.table(paste(main_working_dir, "/HTZ1_sample_id.txt", sep=""), col.names = c("chip", "sample", "strain", "replicate"))
TF="HTZ1"

TF_ID_list$strain <- factor(TF_ID_list$strain)
bam_files = c()
for (i in c(1:dim(TF_ID_list)[1])){
  bam_file_sample=paste(main_working_dir, "/data/elegans/chipseq/alignment/", TF_ID_list[i,1], ".", TF_ID_list[i,3], ".", TF_ID_list[i,4], ".", TF_ID_list[i,2], ".sorted.bam", sep="")
  bam_files = c(bam_files, bam_file_sample)
}

genes <- import.bed(paste(main_working_dir, "/HTZ1/elegans.gene_annotation.coding_gene_longest_isoform.500bp_gene_body.ce11.HTZ1_90pct.bed", sep=""))
genes_tab <- read.table(paste(main_working_dir, "/HTZ1/elegans.gene_annotation.coding_gene_longest_isoform.500bp_gene_body.ce11.HTZ1_90pct.bed", sep=""), col.names = c("chrom", "start", "end", "gene_id"))
bf = BamFileList(bam_files)
se <- summarizeOverlaps(features=genes, bf, mode = "Union")
  
colData(se)$strain <-  factor(TF_ID_list$strain)
de <- DESeqDataSet(se, ~strain)
de1 <- DESeq(de)
  
all_strains=levels(TF_ID_list$strain)
all_mutants=all_strains[all_strains != "N2"]

current_date=gsub("-", "", scan(text = as.character(Sys.time()), what = "")[1])
current_time=gsub(":", "", scan(text = as.character(Sys.time()), what = "")[2])

for (mutant in all_mutants){
  # results against wt
  if ("N2" %in% all_strains){
    res <- results(de1, c('strain', mutant, 'N2'))
    res_genes=as.data.frame(cbind(genes_tab, res))
    downreg=res_genes[res_genes$log2FoldChange < 0 & res_genes$padj < 0.001 & !is.na(res_genes$padj),]
    upreg=res_genes[res_genes$log2FoldChange > 0 & res_genes$padj < 0.001 & !is.na(res_genes$padj),]
    pdf(paste(main_working_dir, "/plots/MA_plots/MA_plot_", mutant, ".", current_date, ".", current_time, ".pdf", sep=""), width=8, height=8)
    plotMA(res, ylim=c(-2,2))
    dev.off()
    write.table(downreg, file = paste(working_dir, TF, "_downregulated_", mutant, "_vs_N2.90_pct.txt", sep=""), quote=F, row.names = F, sep="\t")
    write.table(upreg, file = paste(working_dir, TF, "_upregulated_", mutant, "_vs_N2.90_pct.txt", sep=""), quote=F, row.names = F, sep="\t")
    write.table(res_genes, file = paste(working_dir, TF, "_DESeq_", mutant, "_vs_N2.90_pct.txt", sep=""), quote=F, row.names = F, sep="\t")
}}
```
