## Supplementary material for "DREAM represses distinct targets by cooperating with different THAP domain proteins": Code and related files: DESeq_LIN15BTHAPdel.R

```
library("DESeq2")
library("GenomicAlignments")
library("rtracklayer")


args <- commandArgs()
main_working_dir = args[6]

working_dir=paste(main_working_dir, "/THAPdel_mutants/", sep="")

#Import the peaks and the bam files that I want the counts to be done on 
TF="LIN15B"
strains = c("N2", "lin15BTHAPdel")

TF_ID_list=data.frame("chip" = rep("LIN15B", 6), "sample" = c("AA657", "CG133", "AA1003", "AA999", "CG157", "CG159"), "strain" = c("N2", "N2", "N2", "N2", "lin15BTHAPdel", "lin15BTHAPdel"), "replicate" = c("rep1", "rep2", "rep3", "rep4", "rep1", "rep2"))
TF_ID_list$strain <- factor(TF_ID_list$strain)

bam_files = c()
for (sampleID in TF_ID_list$sample){
  bam_file_sample=grep(sampleID, list.files(path=paste(main_working_dir, "/data/elegans/chipseq/alignment", sep=""), pattern = "\\.sorted\\.bam$", full.names = T), value=T)
  bam_files = c(bam_files, bam_file_sample)
}


peaks <- import.bed(paste(main_working_dir, "/peaks/wt_rescaled/LIN15B_all.peaks.bed", sep=""))
peaks_tab <- read.table(paste(main_working_dir, "/peaks/wt_rescaled/LIN15B_all.peaks.bed", sep=""), col.names = c("chrom", "start", "end", "peak_id"))
bf = BamFileList(bam_files)
se <- summarizeOverlaps(features=peaks, bf, mode = "Union")

colData(se)$strain <-  factor(TF_ID_list$strain)
de <- DESeqDataSet(se, ~strain)
de1 <- DESeq(de)

all_strains=levels(TF_ID_list$strain)
all_mutants=all_strains[all_strains != "N2"]

current_date=gsub("-", "", scan(text = as.character(Sys.time()), what = "")[1])
current_time=gsub(":", "", scan(text = as.character(Sys.time()), what = "")[2])

for (mutant in all_mutants){
  # results against wt
  if ("N2" %in% all_strains){
    res <- results(de1, c('strain', mutant, 'N2'))
    res_peaks=as.data.frame(cbind(peaks_tab, res))
    downreg=res_peaks[res_peaks$log2FoldChange < 0 & res_peaks$padj < 0.01 & !is.na(res_peaks$padj),]
    upreg=res_peaks[res_peaks$log2FoldChange > 0 & res_peaks$padj < 0.01 & !is.na(res_peaks$padj),]
    pdf(paste(main_working_dir, "/plots/MA_plots/MA_plot_", mutant, ".LIN15B_peaks.", current_date, ".", current_time, ".pdf", sep=""), width=8, height=8)
    plotMA(res, ylim=c(-2,2))
    dev.off()
    write.table(downreg, file = paste(working_dir, TF, "_downregulated_", mutant, "_vs_N2.LIN15B_peaks.txt", sep=""), quote=F, row.names = F, sep="\t")
    write.table(upreg, file = paste(working_dir, TF, "_upregulated_", mutant, "_vs_N2.LIN15B_peaks.txt", sep=""), quote=F, row.names = F, sep="\t")
    write.table(res_peaks, file = paste(working_dir, TF, "_DESeq_", mutant, "_vs_N2.LIN15B_peaks.txt", sep=""), quote=F, row.names = F, sep="\t")
  }}
```
