## Supplementary material for "DREAM represses distinct targets by cooperating with different THAP domain proteins": Code and related files: DR_peaks_analysis.R

```
args <- commandArgs()
main_working_dir = args[6]


t_col <- function(color, percent = 50, name = NULL) {
  rgb.val <- col2rgb(color)
  t.col <- rgb(rgb.val[1], rgb.val[2], rgb.val[3],
               max = 255,
               alpha = (100-percent)*255/100,
               names = name)
  invisible(t.col)
  
}

graphical_pvalue = function(x){
  x[x < 0.001] = "***"
  x[x > 0.001 & x < 0.01] = "**"
  x[x > 0.01 & x < 0.05] = "*"
  x[x >= 0.05] = "ns"
  print(x)
}

barplot_data=matrix(rep(0, 36), ncol=3)
barplot_p=matrix(rep(0, 36), ncol=3)

lin_36_DR = read.table(paste(main_working_dir, "/DR_peaks/diff_binding_lin36.hypergeom_test_overlap", sep=""))
lin_36_DR_HT_FC=c()
lin_36_DR_HT_p_value=c()

for (i in c(1:dim(lin_36_DR)[1])){
  lin_36_DR_HT_FC <-  append(lin_36_DR_HT_FC, (lin_36_DR[i,4]/lin_36_DR[i,3]) / (lin_36_DR[i,2]/lin_36_DR[i,1])) #  (x / k ) / (m / N)
  lin_36_DR_HT_p_value <- append(lin_36_DR_HT_p_value, phyper(q=lin_36_DR[i,4] -1, m=lin_36_DR[i,2], n=lin_36_DR[i,1]-lin_36_DR[i,2], k=lin_36_DR[i,3], lower.tail=FALSE))
}

lin_36_DR_fisher_p_value=c()
for (i in seq(1,11,by=2)){
  lin_36_DR_fisher_p_value=append(lin_36_DR_fisher_p_value, fisher.test(matrix(c(lin_36_DR[i,4], lin_36_DR[i,3]-lin_36_DR[i,4], lin_36_DR[i+1,4], lin_36_DR[i+1,3]-lin_36_DR[i+1,4]), ncol=2, byrow=T))$p.value)
}  
lin_36_DR_fisher_p_value_adj = p.adjust(lin_36_DR_fisher_p_value, method="fdr")
lin_36_DR_fisher_p_value_adj_plot=c(lin_36_DR_fisher_p_value_adj[c(1,3,5)], NA, lin_36_DR_fisher_p_value_adj[c(2,4,6)])

lin_36_DR_HT_adj_p_value <- p.adjust(lin_36_DR_HT_p_value, method="fdr")
barplot_data_to_sort = c(lin_36_DR$direct_targets_with_diff_binding/lin_36_DR$n_direct_targets)
barplot_data[,1] = barplot_data_to_sort[c(1,2,5,6,9,10,3,4,7,8,11,12)]
barplot_data[c(2,4,6,8,10,12),1] = 0-barplot_data[c(2,4,6,8,10,12),1]
barplot_p[,1]=lin_36_DR_HT_adj_p_value[c(1,2,5,6,9,10,3,4,7,8,11,12)]
write.table(cbind(row.names(lin_36_DR), lin_36_DR_HT_FC, lin_36_DR_HT_adj_p_value), sep="\t", file = paste(main_working_dir, "/DR_peaks/diff_binding_lin36.hypergeom_test_results", sep=""), quote = F, row.names = F, col.names=F)

lin_35_DR = read.table(paste(main_working_dir, "/DR_peaks/diff_binding_lin35.hypergeom_test_overlap", sep=""))
lin_35_DR_HT_FC=c()
lin_35_DR_HT_p_value=c()

for (i in c(1:dim(lin_35_DR)[1])){
  lin_35_DR_HT_FC <-  append(lin_35_DR_HT_FC, (lin_35_DR[i,4]/lin_35_DR[i,3]) / (lin_35_DR[i,2]/lin_35_DR[i,1])) #  (x / k ) / (m / N)
  lin_35_DR_HT_p_value <- append(lin_35_DR_HT_p_value, phyper(q=lin_35_DR[i,4] -1, m=lin_35_DR[i,2], n=lin_35_DR[i,1]-lin_35_DR[i,2], k=lin_35_DR[i,3], lower.tail=FALSE))
}

lin_35_DR_fisher_p_value=c()
for (i in seq(1,11,by=2)){
  lin_35_DR_fisher_p_value=append(lin_35_DR_fisher_p_value, fisher.test(matrix(c(lin_35_DR[i,4], lin_35_DR[i,3]-lin_35_DR[i,4], lin_35_DR[i+1,4], lin_35_DR[i+1,3]-lin_35_DR[i+1,4]), ncol=2, byrow=T))$p.value)
}  
lin_35_DR_fisher_p_value_adj = p.adjust(lin_35_DR_fisher_p_value, method="fdr")
lin_35_DR_fisher_p_value_adj_plot=c(lin_35_DR_fisher_p_value_adj[c(1,3,5)], NA, lin_35_DR_fisher_p_value_adj[c(2,4,6)])

lin_35_DR_HT_adj_p_value <- p.adjust(lin_35_DR_HT_p_value, method="fdr")
barplot_data_to_sort = c(lin_35_DR$direct_targets_with_diff_binding/lin_35_DR$n_direct_targets)
barplot_data[,2] = barplot_data_to_sort[c(1,2,5,6,9,10,3,4,7,8,11,12)]
barplot_data[c(2,4,6,8,10,12),2] = 0-barplot_data[c(2,4,6,8,10,12),2]
barplot_p[,2]=lin_35_DR_HT_adj_p_value[c(1,2,5,6,9,10,3,4,7,8,11,12)]
write.table(cbind(row.names(lin_35_DR), lin_35_DR_HT_FC, lin_35_DR_HT_adj_p_value), sep="\t", file = paste(main_working_dir, "/DR_peaks/diff_binding_lin35.hypergeom_test_results", sep=""), quote = F, row.names = F, col.names=F)

lin_15B_DR = read.table(paste(main_working_dir, "/DR_peaks/diff_binding_lin15B.hypergeom_test_overlap", sep=""))
lin_15B_DR_HT_FC=c()
lin_15B_DR_HT_p_value=c()

for (i in c(1:dim(lin_15B_DR)[1])){
  lin_15B_DR_HT_FC <-  append(lin_15B_DR_HT_FC, (lin_15B_DR[i,4]/lin_15B_DR[i,3]) / (lin_15B_DR[i,2]/lin_15B_DR[i,1])) #  (x / k ) / (m / N)
  lin_15B_DR_HT_p_value <- append(lin_15B_DR_HT_p_value, phyper(q=lin_15B_DR[i,4] -1, m=lin_15B_DR[i,2], n=lin_15B_DR[i,1]-lin_15B_DR[i,2], k=lin_15B_DR[i,3], lower.tail=FALSE))
}

lin_15B_DR_fisher_p_value=c()
for (i in seq(1,11,by=2)){
  lin_15B_DR_fisher_p_value=append(lin_15B_DR_fisher_p_value, fisher.test(matrix(c(lin_15B_DR[i,4], lin_15B_DR[i,3]-lin_15B_DR[i,4], lin_15B_DR[i+1,4], lin_15B_DR[i+1,3]-lin_15B_DR[i+1,4]), ncol=2, byrow=T))$p.value)
}  
lin_15B_DR_fisher_p_value_adj = p.adjust(lin_15B_DR_fisher_p_value, method="fdr")
lin_15B_DR_fisher_p_value_adj_plot=c(lin_15B_DR_fisher_p_value_adj[c(1,3,5)], NA, lin_15B_DR_fisher_p_value_adj[c(2,4,6)])

lin_15B_DR_HT_adj_p_value <- p.adjust(lin_15B_DR_HT_p_value, method="fdr")
barplot_data_to_sort = c(lin_15B_DR$direct_targets_with_diff_binding/lin_15B_DR$n_direct_targets)
barplot_data[,3] = barplot_data_to_sort[c(1,2,5,6,9,10,3,4,7,8,11,12)]
barplot_data[c(2,4,6,8,10,12),3] = 0-barplot_data[c(2,4,6,8,10,12),3]
barplot_p[,3]=lin_15B_DR_HT_adj_p_value[c(1,2,5,6,9,10,3,4,7,8,11,12)]
write.table(cbind(row.names(lin_15B_DR), lin_15B_DR_HT_FC, lin_15B_DR_HT_adj_p_value), sep="\t", file = paste(main_working_dir, "/DR_peaks/diff_binding_lin15B.hypergeom_test_results", sep=""), quote = F, row.names = F, col.names=F)

barplot_data=rbind(barplot_data[c(1:6),], rep(NA, 6), barplot_data[c(7:12),])
barplot_p=rbind(barplot_p[c(1:6),], rep(NA, 6), barplot_p[c(7:12),])


current_date=gsub("-", "", scan(text = as.character(Sys.time()), what = "")[1])
current_time=gsub(":", "", scan(text = as.character(Sys.time()), what = "")[2])


pdf(paste(main_working_dir, "/plots/ChIP_DR_binding.", current_date, ".", current_time, ".pdf", sep=""), width=7, height=12)
par(mar=c(8,6,4,3), oma=c(0,0,0,0), mfrow=c(3,1))
k=barplot(rep(NA, 7), ylim=c(-1,1), axes=F) 
abline(h=c(0, 0.5, -0.5), lty=2, lwd=2, col=c("grey40", t_col("grey", 50), t_col("grey", 50)), )
k=barplot(barplot_data[c(1,3,5,7,8,10,12),1], ylim=c(-1,1), beside=T, las=1, add=T, col=t_col("forestgreen", 50), ylab="up/downregulated peak fraction", main = "binding on direct targets in lin-36") 
k=barplot(barplot_data[c(2,4,6,7,9,11,13),1], ylim=c(-1,1), beside=T, las=1, add=T, col=t_col("indianred", 50)) 
text(k[,1], 1, graphical_pvalue(lin_36_DR_fisher_p_value_adj_plot), pos = 1)
text(k[c(1,2,3,5,6,7),1], -1.15, labels = rep(c("LIN-35", "EFL-1", "LIN-15B"), 2), xpd=T)
segments(c(k[1,1], k[5,1]), -1.25, c(k[3,1], k[7,1]), -1.25, xpd=T)
text(c(k[2,1], k[6,1]), -1.35, labels = c("LIN-36 shared", "LIN-15B shared"), xpd=T)

k=barplot(rep(NA, 7), ylim=c(-1,1), axes=F) 
abline(h=c(0, 0.5, -0.5), lty=2, lwd=2, col=c("grey40", t_col("grey", 50), t_col("grey", 50)), )
k=barplot(barplot_data[c(1,3,5,7,8,10,12),2], ylim=c(-1,1), beside=T, las=1, add=T, col=t_col("forestgreen", 50), ylab="up/downregulated peak fraction", main = "binding on direct targets in lin-35") 
k=barplot(barplot_data[c(2,4,6,7,9,11,13),2], ylim=c(-1,1), beside=T, las=1, add=T, col=t_col("indianred", 50)) 
text(k[,1], 1, graphical_pvalue(lin_35_DR_fisher_p_value_adj_plot), pos = 1)
text(k[c(1,2,3,5,6,7),1], -1.15, labels = rep(c("EFL-1", "LIN-36", "LIN-15B"), 2), xpd=T)
segments(c(k[1,1], k[5,1]), -1.25, c(k[3,1], k[7,1]), -1.25, xpd=T)
text(c(k[2,1], k[6,1]), -1.35, labels = c("LIN-36 shared", "LIN-15B shared"), xpd=T)

k=barplot(rep(NA, 7), ylim=c(-1,1), axes=F) 
abline(h=c(0, 0.5, -0.5), lty=2, lwd=2, col=c("grey40", t_col("grey", 50), t_col("grey", 50)), )
k=barplot(barplot_data[c(1,3,5,7,8,10,12),3], ylim=c(-1,1), beside=T, las=1, add=T, col=t_col("forestgreen", 50), ylab="up/downregulated peak fraction", main = "binding on direct targets in lin-15B") 
k=barplot(barplot_data[c(2,4,6,7,9,11,13),3], ylim=c(-1,1), beside=T, las=1, add=T, col=t_col("indianred", 50)) 
text(k[,1], 1, graphical_pvalue(lin_15B_DR_fisher_p_value_adj_plot), pos = 1)
text(k[c(1,2,3,5,6,7),1], -1.15, labels = rep(c("LIN-35", "EFL-1", "LIN-36"), 2), xpd=T)
segments(c(k[1,1], k[5,1]), -1.25, c(k[3,1], k[7,1]), -1.25, xpd=T)
text(c(k[2,1], k[6,1]), -1.35, labels = c("LIN-36 shared", "LIN-15B shared"), xpd=T)
dev.off()


barplot_data=matrix(rep(0, 4), ncol=1)
barplot_p=matrix(rep(0, 4), ncol=1)


lin_15BTHAPdel_DR = read.table(paste(main_working_dir, "/DR_peaks/diff_binding_lin15BTHAPdel.hypergeom_test_overlap", sep=""))
lin_15BTHAPdel_DR_HT_FC=c()
lin_15BTHAPdel_DR_HT_p_value=c()

for (i in c(1:dim(lin_15BTHAPdel_DR)[1])){
  lin_15BTHAPdel_DR_HT_FC <-  append(lin_15BTHAPdel_DR_HT_FC, (lin_15BTHAPdel_DR[i,4]/lin_15BTHAPdel_DR[i,3]) / (lin_15BTHAPdel_DR[i,2]/lin_15BTHAPdel_DR[i,1])) #  (x / k ) / (m / N)
  lin_15BTHAPdel_DR_HT_p_value <- append(lin_15BTHAPdel_DR_HT_p_value, phyper(q=lin_15BTHAPdel_DR[i,4] -1, m=lin_15BTHAPdel_DR[i,2], n=lin_15BTHAPdel_DR[i,1]-lin_15BTHAPdel_DR[i,2], k=lin_15BTHAPdel_DR[i,3], lower.tail=FALSE))
}

lin_15BTHAPdel_DR_fisher_p_value=c()
for (i in seq(1,3,by=2)){
  lin_15BTHAPdel_DR_fisher_p_value=append(lin_15BTHAPdel_DR_fisher_p_value, fisher.test(matrix(c(lin_15BTHAPdel_DR[i,4], lin_15BTHAPdel_DR[i,3]-lin_15BTHAPdel_DR[i,4], lin_15BTHAPdel_DR[i+1,4], lin_15BTHAPdel_DR[i+1,3]-lin_15BTHAPdel_DR[i+1,4]), ncol=2, byrow=T))$p.value)
}
lin_15BTHAPdel_DR_fisher_p_value_adj = p.adjust(lin_15BTHAPdel_DR_fisher_p_value, method="fdr")
lin_15BTHAPdel_DR_fisher_p_value_adj_plot=c(lin_15BTHAPdel_DR_fisher_p_value_adj[1], NA, lin_15BTHAPdel_DR_fisher_p_value_adj[2])

lin_15BTHAPdel_DR_HT_adj_p_value <- p.adjust(lin_15BTHAPdel_DR_HT_p_value, method="fdr")
barplot_data_to_sort = c(lin_15BTHAPdel_DR$direct_targets_with_diff_binding/lin_15BTHAPdel_DR$n_direct_targets)
barplot_data[,1] = barplot_data_to_sort[c(1,2,3,4)]
barplot_data[c(2,4),1] = 0-barplot_data[c(2,4),1]
barplot_p[,1]=lin_15BTHAPdel_DR_HT_adj_p_value[c(1,2,3,4)]
write.table(cbind(row.names(lin_15BTHAPdel_DR), lin_15BTHAPdel_DR_HT_FC, lin_15BTHAPdel_DR_HT_adj_p_value), sep="\t", file = paste(main_working_dir, "/DR_peaks/diff_binding_lin15BTHAPdel.hypergeom_test_results", sep=""), quote = F, row.names = F, col.names=F)

barplot_data=c(barplot_data[c(1:2),], NA, barplot_data[c(3:4),])
barplot_p=c(barplot_p[c(1:2),], NA, barplot_p[c(3:4),])


current_date=gsub("-", "", scan(text = as.character(Sys.time()), what = "")[1])
current_time=gsub(":", "", scan(text = as.character(Sys.time()), what = "")[2])


pdf(paste(main_working_dir, "/plots/ChIP_DR_binding.LIN15BTHAPdel.", current_date, ".", current_time, ".pdf", sep=""), width=7, height=12)
par(mar=c(8,6,4,3), oma=c(0,0,0,0), mfrow=c(1,1))
k=barplot(rep(NA, 3), ylim=c(-1,1), axes=F)
abline(h=c(0, 0.5, -0.5), lty=2, lwd=2, col=c("grey40", t_col("grey", 50), t_col("grey", 50)), )
k=barplot(barplot_data[c(1,3,4)], ylim=c(-1,1), beside=T, las=1, add=T, col=t_col("forestgreen", 50), ylab="up/downregulated peak fraction", main = "binding on direct targets in lin-15BTHAPdel")
k=barplot(barplot_data[c(2,3,5)], ylim=c(-1,1), beside=T, las=1, add=T, col=t_col("indianred", 50))
text(k[,1], 1, graphical_pvalue(lin_15BTHAPdel_DR_fisher_p_value_adj_plot), pos = 1)
text(k[2], -1.15, labels = "LIN-15BTHAPdel", xpd=T)
dev.off()
```
