## Supplementary material for "DREAM represses distinct targets by cooperating with different THAP domain proteins": Code and related files: E2F_strength.R

```
# MEME E2F motifs enrichment
args <- commandArgs()
main_working_dir = args[6]

t_col <- function(color, percent = 50, name = NULL) {
  rgb.val <- col2rgb(color)
  t.col <- rgb(rgb.val[1], rgb.val[2], rgb.val[3],
               max = 255,
               alpha = (100-percent)*255/100,
               names = name)
  invisible(t.col)
}

graphical_pvalue = function(x){
  x[x < 0.001] = "***"
  x[x > 0.001 & x < 0.01] = "**"
  x[x > 0.01 & x < 0.05] = "*"
  x[x >= 0.05] = "ns"
  print(x)
}

current_date=gsub("-", "", scan(text = as.character(Sys.time()), what = "")[1])
current_time=gsub(":", "", scan(text = as.character(Sys.time()), what = "")[2])

lin36_shared_col=t_col("lightgreen", 50)
lin15B_shared_col=t_col("mediumpurple1", 50)

LIN36_shared_LIN15B_motif = read.table(paste(main_working_dir, "/motif_analysis/E2F_enrichment/fimo_LIN35_LIN36_shared.motif_LIN35_LIN15B/fimo.max_score.bed", sep=""))
LIN15B_shared_LIN15B_motif = read.table(paste(main_working_dir, "/motif_analysis/E2F_enrichment/fimo_LIN35_LIN15B_shared.motif_LIN35_LIN15B/fimo.max_score.bed", sep=""))
LIN36_shared_LIN36_motif = read.table(paste(main_working_dir, "/motif_analysis/E2F_enrichment/fimo_LIN35_LIN36_shared.motif_LIN35_LIN36/fimo.max_score.bed", sep=""))
LIN15B_shared_LIN36_motif = read.table(paste(main_working_dir, "/motif_analysis/E2F_enrichment/fimo_LIN35_LIN15B_shared.motif_LIN35_LIN36/fimo.max_score.bed", sep=""))

LIN36_shared_peaks = read.table(paste(main_working_dir, "/direct_target_peaks_promoters/LIN35_LIN36GFP_shared.lin35_lin36full_direct_targets.bed", sep=""))
LIN15B_shared_peaks = read.table(paste(main_working_dir, "/direct_target_peaks_promoters/LIN35_LIN15B_shared.lin35_lin15B_direct_targets.bed", sep=""))

LIN15B_motif_abundance=matrix(c(dim(LIN36_shared_LIN15B_motif[LIN36_shared_LIN15B_motif$V5 < 0.001,])[1],
                                dim(LIN36_shared_peaks)[1]-dim(LIN36_shared_LIN15B_motif[LIN36_shared_LIN15B_motif$V5 < 0.001,])[1],
                                dim(LIN15B_shared_LIN15B_motif[LIN15B_shared_LIN15B_motif$V5 < 0.001,])[1],
                                dim(LIN15B_shared_peaks)[1]-dim(LIN15B_shared_LIN15B_motif[LIN15B_shared_LIN15B_motif$V5 < 0.001,])[1]), ncol=2, byrow=T)
fisher.test(LIN15B_motif_abundance)

LIN36_motif_abundance=matrix(c(dim(LIN36_shared_LIN36_motif[LIN36_shared_LIN36_motif$V5 < 0.001,])[1],
                               dim(LIN36_shared_peaks)[1]-dim(LIN36_shared_LIN36_motif[LIN36_shared_LIN36_motif$V5 < 0.001,])[1],
                               dim(LIN15B_shared_LIN36_motif[LIN15B_shared_LIN36_motif$V5 < 0.001,])[1],
                               dim(LIN15B_shared_peaks)[1]-dim(LIN15B_shared_LIN36_motif[LIN15B_shared_LIN36_motif$V5 < 0.001,])[1]), ncol=2, byrow=T)
fisher.test(LIN36_motif_abundance)

pdf(paste(main_working_dir, "/plots/E2F_abundance.", current_date, ".", current_time, ".pdf", sep=""), width=6, height=6)

par(mfrow=c(1,1), mar=c(3,5,3,3), oma=c(2,0,0,0))
k=barplot(c(LIN36_motif_abundance[,1]/rowSums(LIN36_motif_abundance), LIN15B_motif_abundance[,1]/rowSums(LIN15B_motif_abundance)), ylim=c(0, 1), col=c(lin36_shared_col, lin15B_shared_col), las=1, ylab="fraction of promoters", main = "strong E2F motif abundance")
axis(1, at=c((k[1,1]+k[2,1])/2,(k[3,1]+k[4,1])/2), labels = c("E2F-a", "E2F-b"), line = 0.5) 
segments(c(k[1,1], k[3,1]), 0.9, c(k[2,1], k[4,1]), 0.9)
text(c((k[1,1]+k[2,1])/2,(k[3,1]+k[4,1])/2), 0.9, labels = graphical_pvalue(c(fisher.test(LIN36_motif_abundance)$p.value,
                                                                            fisher.test(LIN15B_motif_abundance)$p.value)), pos=3)
par(fig = c(0, 1, 0, 1), oma = c(0, 0, 0, 0), mar = c(0, 0, 0, 0), new = TRUE)
plot(0, 0, type = "n", bty = "n", xaxt = "n", yaxt = "n")
legend(0, -0.95, legend = c("LIN-36-shared", "LIN-15B-shared"), fill = c(lin36_shared_col, lin15B_shared_col), bty="n", ncol = 2, xjust = 0.2)
dev.off()

wilcox.test(LIN15B_shared_LIN36_motif$V4,LIN36_shared_LIN36_motif$V4)
wilcox.test(LIN15B_shared_LIN15B_motif$V4,LIN36_shared_LIN15B_motif$V4)

pdf(paste(main_working_dir, "/plots/E2F_strength.", current_date, ".", current_time, ".pdf", sep=""), width=6, height=6)

par(mfrow=c(1,1), mar=c(3,5,3,3), oma=c(2,0,0,0))
x<-boxplot(LIN36_shared_LIN36_motif$V4,
           LIN15B_shared_LIN36_motif$V4,
           LIN36_shared_LIN15B_motif$V4,
           LIN15B_shared_LIN15B_motif$V4, plot=F)
ylim_plot=c(min(c(x$stats, x$out)), max(c(x$stats, x$out)))
boxplot(LIN36_shared_LIN36_motif$V4,
        LIN15B_shared_LIN36_motif$V4,
        LIN36_shared_LIN15B_motif$V4,
        LIN15B_shared_LIN15B_motif$V4, notch=T, las=1, 
        ylim=c(ylim_plot[1], ylim_plot[2]+(ylim_plot[2]-ylim_plot[1])/15), 
        xaxt="n", main = "E2F strength", ylab = "motif strength", col=c(lin36_shared_col, lin15B_shared_col))
axis(1, at=c(1.5,3.5), labels = c("E2F-a", "E2F-b"))
segments(c(1,3), ylim_plot[2]+(ylim_plot[2]-ylim_plot[1])/20, c(2,4), ylim_plot[2]+(ylim_plot[2]-ylim_plot[1])/20)
text(c(1.5,3.5), ylim_plot[2]+(ylim_plot[2]-ylim_plot[1])/20, labels = graphical_pvalue(c(wilcox.test(LIN15B_shared_LIN36_motif$V4,
                                                                                                      LIN36_shared_LIN36_motif$V4)$p.value,
                                                                                          wilcox.test(LIN15B_shared_LIN15B_motif$V4,
                                                                                                      LIN36_shared_LIN15B_motif$V4)$p.value)), pos=3)
par(fig = c(0, 1, 0, 1), oma = c(0, 0, 0, 0), mar = c(0, 0, 0, 0), new = TRUE)
plot(0, 0, type = "n", bty = "n", xaxt = "n", yaxt = "n")
legend(0, -0.95, legend = c("LIN-36-shared", "LIN-15B-shared"), fill = c(lin36_shared_col, lin15B_shared_col), bty="n", ncol = 2, xjust = 0.2)
dev.off()
```
