## Supplementary material for "DREAM represses distinct targets by cooperating with different THAP domain proteins": Code and related files: gene_biotype_def.py

```
import sys

gff_in=sys.argv[1]
gene_annotation=sys.argv[2]
gene_transcript=sys.argv[3]
biotype_table_out=sys.argv[4]
coding_genes_out=sys.argv[5]

new = open(biotype_table_out, "w")
coding_genes=[]

for line in open(gff_in):
    G=line.rstrip().split("\t")
    if G[2] == "gene":
        biotype=""
        K=G[8].split(";")
        for i in K:
            if "ID=" in i:
                if ":" in i:
                    gene = i.split(":")[1]
                else:
                    gene = i.split("=")[1]
            if "biotype" in i:
                biotype=i.split("=")[1]
        if biotype == "":
            biotype="protein_coding"
        temp = gene + "\t" + biotype + "\n"
        new.write(temp)
        if biotype== "protein_coding":
            coding_genes.append(gene)

new.close()

coding_tr=[]
for line in open(gene_transcript):
    G=line.rstrip().split("\t")
    if G[0] in coding_genes:
        coding_tr.append(G[1])

new = open(coding_genes_out, "w")
for line in open(gene_annotation):
    G=line.rstrip().split("\t")
    if G[3] in coding_tr:
        new.write(line)

new.close()
```
