## Supplementary material for "DREAM represses distinct targets by cooperating with different THAP domain proteins": Code and related files: gene_covarage.sh

```
#!/bin/bash

working_dir=$1
sample_name=$2
strain_name=$3

coverageBed -a $working_dir/species/elegans/gene_annotation/elegans.gene_annotation.coding_gene_longest_isoform.ce11.bed -b $working_dir/data/elegans/RNASeq/alignment/$strain_name/$strain_name.$sample_name.Aligned.sortedByCoord.out.bam -split | cut -f 4,13,14,15,16 > $working_dir/data/elegans/RNASeq/alignment/$strain_name/$strain_name.$sample_name.gene_coverage.txt
```
