## Supplementary material for "DREAM represses distinct targets by cooperating with different THAP domain proteins": Code and related files: gene_longest_isoform_annotation.py

```
import sys

transcript_file=sys.argv[1]
transcript_gene=sys.argv[2]
output_file=sys.argv[3]
species=sys.argv[4]

tr_gene={}
for line in open(transcript_gene):
    G=line.rstrip().split("\t")
    tr_gene[G[1]] = G[0]

longest_isoform = {}
for line in open(transcript_file):
    G=line.rstrip().split("\t")
    if tr_gene[G[3]] not in longest_isoform:
        longest_isoform[tr_gene[G[3]]] = line
    else:
        K=G[10].split(",")
        if sum(int(y) for y in K[:-1]) > sum(int(x) for x in longest_isoform[tr_gene[G[3]]].split("\t")[10].split(",")[:-1]):
            longest_isoform[tr_gene[G[3]]] = line

new=open(output_file+".bed", "w")
for gene in longest_isoform:
    new_gene_name = gene
    K = longest_isoform[gene].rstrip().split("\t")
    temp=""
    K[3] = new_gene_name
    for i in K:
        temp += i + "\t"
    new.write(temp.rstrip() + "\n")

new.close()

new=open(output_file+".longest_isoform", "w")
for gene in longest_isoform:
    new_gene_name = gene
    K = longest_isoform[gene].rstrip().split("\t")
    new.write(K[3] + "\t" + new_gene_name + "\n")

new.close()
```
