## Supplementary material for "DREAM represses distinct targets by cooperating with different THAP domain proteins": Code and related files: gene_longest_isoform_annotation.sh

```
#!/bin/bash

species=$1
scripts_path=$2
species_path=$3

python $scripts_path/gene_longest_isoform_annotation.py $species_path"/"$species"/gene_annotation/"$species".gene_annotation.coding_gene.ce11.bed" $species_path"/"$species"/gene_annotation/"$species".gene_annotation.gene_transcript_id.ce11.txt" $species_path"/"$species"/gene_annotation/"$species".gene_annotation.coding_gene_longest_isoform.ce11" $species
```
