## Supplementary material for "DREAM represses distinct targets by cooperating with different THAP domain proteins": Code and related files: gene_transcript_id_extractor.py

```
import sys

gff_in=sys.argv[1]
table_out=sys.argv[2]

new = open(table_out, "w")

for line in open(gff_in):
    G=line.rstrip().split("\t")
    if G[2] == "mRNA":
        K = G[8].split(";")
        if len(K) == 1:
            if K[0][:3] == "ID=":
                transcript=K[0].split("=")[1]
                gene=K[0].split("=")[1]
        else:
            for i in K:
                if i[:3] == "ID=":
                    if ":" in i:
                        transcript=i.split(":")[1]
                    else:
                        transcript=i.split("=")[1]
                if i[:7] == "Parent=":
                    if ":" in i:
                        gene=i.split(":")[1]
                    else:
                        gene=i.split("=")[1]
        temp = gene + "\t" + transcript + "\n"
        new.write(temp)

new.close()
```
