## Supplementary material for "DREAM represses distinct targets by cooperating with different THAP domain proteins": Code and related files: H3K9me2_direct_target_peaks_coverage.R

```
# compare H3K9me2 coverage over gene body of LIN15B-shared and LIN36-shared direct targets
library("beeswarm")

args <- commandArgs()
main_working_dir = args[6]

t_col <- function(color, percent = 50, name = NULL) {
  rgb.val <- col2rgb(color)
  t.col <- rgb(rgb.val[1], rgb.val[2], rgb.val[3],
               max = 255,
               alpha = (100-percent)*255/100,
               names = name)
  invisible(t.col)
  
}

graphical_pvalue = function(x){
  x[x < 0.001] = "***"
  x[x > 0.001 & x < 0.01] = "**"
  x[x > 0.01 & x < 0.05] = "*"
  x[x >= 0.05] = "ns"
  print(x)
}

lin35_only_col=t_col("coral2", 50)
lin36_shared_col=t_col("lightgreen", 50)
lin15B_shared_col=t_col("mediumpurple1", 50)


lin15B_shared_H3K9me2_N2 = read.table(paste(main_working_dir, "/H3K9me2/LIN35_LIN15B_shared.lin35_lin15B_direct_target_peaks.H3K9me2_N2.bed", sep=""))
lin36_shared_H3K9me2_N2 = read.table(paste(main_working_dir, "/H3K9me2/LIN35_LIN36GFP_shared.lin35_lin36full_direct_target_peaks.H3K9me2_N2.bed", sep=""))
lin35_only_H3K9me2_N2 = read.table(paste(main_working_dir, "/H3K9me2/LIN35_only.lin35_direct_target_peaks.H3K9me2_N2.bed", sep=""))

lin15B_lin36_P=wilcox.test(lin36_shared_H3K9me2_N2$V4, lin15B_shared_H3K9me2_N2$V4)$p.value

current_date=gsub("-", "", scan(text = as.character(Sys.time()), what = "")[1])
current_time=gsub(":", "", scan(text = as.character(Sys.time()), what = "")[2])

pdf(paste(main_working_dir, "/plots/H3K9me2_direct_targets_coverage.N2.", current_date, ".", current_time, ".pdf", sep=""), width=5, height=5)
par(mfrow=c(1,1), mar=c(5,5,3,3), oma=c(0,0,0,0))
x<-boxplot(lin36_shared_H3K9me2_N2$V4,
           lin15B_shared_H3K9me2_N2$V4, plot=F)
ylim_plot=c(min(c(x$stats, x$out)), max(c(x$stats, x$out)))
x<-boxplot(lin36_shared_H3K9me2_N2$V4,
           lin15B_shared_H3K9me2_N2$V4, notch=T, las=1, xaxt="n", ylim=c(ylim_plot[1], ylim_plot[2]+(ylim_plot[2]-ylim_plot[1])/15),
           ylab="H3K9me2 coverage", main="direct target promoters H3K9me2", col=c(lin36_shared_col, lin15B_shared_col), outline=F)
text(cex=1, x=c(2,3), y=ylim_plot[1]-(ylim_plot[2]-ylim_plot[1])/15, c("LIN-36-shared", "LIN-15B-shared"), xpd=TRUE, srt=30, adj = 1)
beeswarm(list(lin36_shared_H3K9me2_N2$V4,
              lin15B_shared_H3K9me2_N2$V4), add=T, at = c(1,2),
         vertical = T, method = "swarm", spacing=1, corral="wrap", pch = 16, col = t_col("grey60", 50), cex = 0.6)
text(cex=1, 1.5, y=ylim_plot[2]-(ylim_plot[2]-ylim_plot[1])/10, graphical_pvalue(lin15B_lin36_P), adj = c(0, 0))
segments(1, ylim_plot[2]-(ylim_plot[2]-ylim_plot[1])/10, 
         2, ylim_plot[2]-(ylim_plot[2]-ylim_plot[1])/10)
	 
dev.off()


# number of peaks with lower H3K9me2 levels in mutants
lin35_H3K9me2_down = read.table(paste(main_working_dir, "/H3K9me2/H3K9me2_downregulated_lin35_vs_N2.H3K9me2_peaks.txt", sep=""), header = T)
lin36_H3K9me2_down = read.table(paste(main_working_dir, "/H3K9me2/H3K9me2_downregulated_lin36full_vs_N2.H3K9me2_peaks.txt", sep=""), header = T)
lin15B_H3K9me2_down = read.table(paste(main_working_dir, "/H3K9me2/H3K9me2_downregulated_lin15B_vs_N2.H3K9me2_peaks.txt", sep=""), header = T)

pdf(paste(main_working_dir, "/plots/H3K9me2_downreg_peaks_mutants.", current_date, ".", current_time, ".pdf", sep=""), width=6, height=6)
par(mfrow=c(1,1), mar=c(3,5,3,3), oma=c(0,0,0,0))
barplot(c(dim(lin35_H3K9me2_down)[1],
          dim(lin36_H3K9me2_down)[1],
          dim(lin15B_H3K9me2_down)[1]), beside=T, las=1, 
        col=c(lin35_only_col, lin36_shared_col, lin15B_shared_col), names.arg = c(expression(italic("lin-35")), expression(italic("lin-36")), expression(italic("lin-15B"))), ylab="nº of peaks", main = "H3K9me2 downregulated peaks")
dev.off()
```
