## Supplementary material for "DREAM represses distinct targets by cooperating with different THAP domain proteins": Code and related files: H3K9me2_loss_hypergeom_test.R

```
args <- commandArgs()
main_working_dir = args[6]


t_col <- function(color, percent = 50, name = NULL) {
  rgb.val <- col2rgb(color)
  t.col <- rgb(rgb.val[1], rgb.val[2], rgb.val[3],
               max = 255,
               alpha = (100-percent)*255/100,
               names = name)
  invisible(t.col)
}

graphical_pvalue = function(x){
  x[x < 0.001] = "***"
  x[x > 0.001 & x < 0.01] = "**"
  x[x > 0.01 & x < 0.05] = "*"
  x[x >= 0.05] = "ns"
  print(x)
}

barplot_data=matrix(rep(0, 24), ncol=3)

lin_36_shared_H3K9me2 = read.table(paste(main_working_dir, "/H3K9me2/LIN35_LIN36GFP_shared.lin35_lin36full.hypergeom_test_overlap", sep=""))
lin_36_shared_HT_FC=c()
lin_36_shared_HT_p_value=c()

for (i in c(1:dim(lin_36_shared_H3K9me2)[1])){
  lin_36_shared_HT_FC <-  append(lin_36_shared_HT_FC, (lin_36_shared_H3K9me2[i,4]/lin_36_shared_H3K9me2[i,3]) / (lin_36_shared_H3K9me2[i,2]/lin_36_shared_H3K9me2[i,1])) #  (x / k ) / (m / N)
  lin_36_shared_HT_p_value <- append(lin_36_shared_HT_p_value, phyper(q=lin_36_shared_H3K9me2[i,4] -1, m=lin_36_shared_H3K9me2[i,2], n=lin_36_shared_H3K9me2[i,1]-lin_36_shared_H3K9me2[i,2], k=lin_36_shared_H3K9me2[i,3], lower.tail=FALSE))
}

lin_36_shared_HT_adj_p_value <- p.adjust(lin_36_shared_HT_p_value, method="fdr")
barplot_data[,1] = c(lin_36_shared_H3K9me2$direct_targets_with_low_H3K9me2, lin_36_shared_H3K9me2$n_direct_targets[1]-sum(lin_36_shared_H3K9me2$direct_targets_with_low_H3K9me2))
write.table(cbind(row.names(lin_36_shared_H3K9me2), lin_36_shared_HT_FC, lin_36_shared_HT_adj_p_value), sep="\t", file = paste(main_working_dir, "/H3K9me2/LIN35_LIN36GFP_shared.lin35_lin36full.hypergeom_test_results", sep=""), quote = F, row.names = F, col.names=F)

lin_36_shared_H3K9me2_by_factor=data.frame(row.names=c("lin35", "lin36", "lin15B"), "tested_peaks" = rep(lin_36_shared_H3K9me2[1,1], 3), "low_H3K9me2_in_mutant" = c(lin_36_shared_H3K9me2[1,2]+lin_36_shared_H3K9me2[4,2]+lin_36_shared_H3K9me2[5,2]+lin_36_shared_H3K9me2[7,2],
                                                                                                                                                              lin_36_shared_H3K9me2[2,2]+lin_36_shared_H3K9me2[4,2]+lin_36_shared_H3K9me2[6,2]+lin_36_shared_H3K9me2[7,2],
                                                                                                                                                              lin_36_shared_H3K9me2[3,2]+lin_36_shared_H3K9me2[5,2]+lin_36_shared_H3K9me2[6,2]+lin_36_shared_H3K9me2[7,2]),
                                        "n_direct_targets" = rep(lin_36_shared_H3K9me2[1,3], 3), "direct_targets_with_low_H3K9me2" = c(lin_36_shared_H3K9me2[1,4]+lin_36_shared_H3K9me2[4,4]+lin_36_shared_H3K9me2[5,4]+lin_36_shared_H3K9me2[7,4],
                                                                                                                                   lin_36_shared_H3K9me2[2,4]+lin_36_shared_H3K9me2[4,4]+lin_36_shared_H3K9me2[6,4]+lin_36_shared_H3K9me2[7,4],
                                                                                                                                   lin_36_shared_H3K9me2[3,4]+lin_36_shared_H3K9me2[5,4]+lin_36_shared_H3K9me2[6,4]+lin_36_shared_H3K9me2[7,4]))
lin_36_shared_by_factor_HT_FC=c()
lin_36_shared_by_factor_HT_p_value=c()

for (i in c(1:dim(lin_36_shared_H3K9me2_by_factor)[1])){
  lin_36_shared_by_factor_HT_FC <-  append(lin_36_shared_by_factor_HT_FC, (lin_36_shared_H3K9me2_by_factor[i,4]/lin_36_shared_H3K9me2_by_factor[i,3]) / (lin_36_shared_H3K9me2_by_factor[i,2]/lin_36_shared_H3K9me2_by_factor[i,1])) #  (x / k ) / (m / N)
  lin_36_shared_by_factor_HT_p_value <- append(lin_36_shared_by_factor_HT_p_value, phyper(q=lin_36_shared_H3K9me2_by_factor[i,4] -1, m=lin_36_shared_H3K9me2_by_factor[i,2], n=lin_36_shared_H3K9me2_by_factor[i,1]-lin_36_shared_H3K9me2_by_factor[i,2], k=lin_36_shared_H3K9me2_by_factor[i,3], lower.tail=FALSE))
}
lin_36_shared_by_factor_HT_adj_p_value=p.adjust(lin_36_shared_by_factor_HT_p_value, method="fdr")


lin_15B_shared_H3K9me2 = read.table(paste(main_working_dir, "/H3K9me2/LIN35_LIN15B_shared.lin35_lin15B.hypergeom_test_overlap", sep=""))
lin_15B_shared_HT_FC=c()
lin_15B_shared_HT_p_value=c()

for (i in c(1:dim(lin_15B_shared_H3K9me2)[1])){
  lin_15B_shared_HT_FC <-  append(lin_15B_shared_HT_FC, (lin_15B_shared_H3K9me2[i,4]/lin_15B_shared_H3K9me2[i,3]) / (lin_15B_shared_H3K9me2[i,2]/lin_15B_shared_H3K9me2[i,1])) #  (x / k ) / (m / N)
  lin_15B_shared_HT_p_value <- append(lin_15B_shared_HT_p_value, phyper(q=lin_15B_shared_H3K9me2[i,4] -1, m=lin_15B_shared_H3K9me2[i,2], n=lin_15B_shared_H3K9me2[i,1]-lin_15B_shared_H3K9me2[i,2], k=lin_15B_shared_H3K9me2[i,3], lower.tail=FALSE))
}

lin_15B_shared_HT_adj_p_value <- p.adjust(lin_15B_shared_HT_p_value, method="fdr")
barplot_data[,2] = c(lin_15B_shared_H3K9me2$direct_targets_with_low_H3K9me2, lin_15B_shared_H3K9me2$n_direct_targets[1]-sum(lin_15B_shared_H3K9me2$direct_targets_with_low_H3K9me2))
write.table(cbind(row.names(lin_15B_shared_H3K9me2), lin_15B_shared_HT_FC, lin_15B_shared_HT_adj_p_value), sep="\t", file = paste(main_working_dir, "/H3K9me2/LIN35_LIN15B_shared.lin35_lin15B.hypergeom_test_results", sep=""), quote = F, row.names = F, col.names=F)

lin_15B_shared_H3K9me2_by_factor=data.frame(row.names=c("lin35", "lin36", "lin15B"), "tested_peaks" = rep(lin_15B_shared_H3K9me2[1,1], 3), "low_H3K9me2_in_mutant" = c(lin_15B_shared_H3K9me2[1,2]+lin_15B_shared_H3K9me2[4,2]+lin_15B_shared_H3K9me2[5,2]+lin_15B_shared_H3K9me2[7,2],
                                                                                                                                                                     lin_15B_shared_H3K9me2[2,2]+lin_15B_shared_H3K9me2[4,2]+lin_15B_shared_H3K9me2[6,2]+lin_15B_shared_H3K9me2[7,2],
                                                                                                                                                                     lin_15B_shared_H3K9me2[3,2]+lin_15B_shared_H3K9me2[5,2]+lin_15B_shared_H3K9me2[6,2]+lin_15B_shared_H3K9me2[7,2]),
                                           "n_direct_targets" = rep(lin_15B_shared_H3K9me2[1,3], 3), "direct_targets_with_low_H3K9me2" = c(lin_15B_shared_H3K9me2[1,4]+lin_15B_shared_H3K9me2[4,4]+lin_15B_shared_H3K9me2[5,4]+lin_15B_shared_H3K9me2[7,4],
                                                                                                                                          lin_15B_shared_H3K9me2[2,4]+lin_15B_shared_H3K9me2[4,4]+lin_15B_shared_H3K9me2[6,4]+lin_15B_shared_H3K9me2[7,4],
                                                                                                                                          lin_15B_shared_H3K9me2[3,4]+lin_15B_shared_H3K9me2[5,4]+lin_15B_shared_H3K9me2[6,4]+lin_15B_shared_H3K9me2[7,4]))
lin_15B_shared_by_factor_HT_FC=c()
lin_15B_shared_by_factor_HT_p_value=c()

for (i in c(1:dim(lin_15B_shared_H3K9me2_by_factor)[1])){
  lin_15B_shared_by_factor_HT_FC <-  append(lin_15B_shared_by_factor_HT_FC, (lin_15B_shared_H3K9me2_by_factor[i,4]/lin_15B_shared_H3K9me2_by_factor[i,3]) / (lin_15B_shared_H3K9me2_by_factor[i,2]/lin_15B_shared_H3K9me2_by_factor[i,1])) #  (x / k ) / (m / N)
  lin_15B_shared_by_factor_HT_p_value <- append(lin_15B_shared_by_factor_HT_p_value, phyper(q=lin_15B_shared_H3K9me2_by_factor[i,4] -1, m=lin_15B_shared_H3K9me2_by_factor[i,2], n=lin_15B_shared_H3K9me2_by_factor[i,1]-lin_15B_shared_H3K9me2_by_factor[i,2], k=lin_15B_shared_H3K9me2_by_factor[i,3], lower.tail=FALSE))
}
lin_15B_shared_by_factor_HT_adj_p_value=p.adjust(lin_15B_shared_by_factor_HT_p_value, method="fdr")


lin_35_only_H3K9me2 = read.table(paste(main_working_dir, "/H3K9me2/LIN35_only.lin35.hypergeom_test_overlap", sep=""))
lin_35_only_HT_FC=c()
lin_35_only_HT_p_value=c()

for (i in c(1:dim(lin_35_only_H3K9me2)[1])){
  lin_35_only_HT_FC <-  append(lin_35_only_HT_FC, (lin_35_only_H3K9me2[i,4]/lin_35_only_H3K9me2[i,3]) / (lin_35_only_H3K9me2[i,2]/lin_35_only_H3K9me2[i,1])) #  (x / k ) / (m / N)
  lin_35_only_HT_p_value <- append(lin_35_only_HT_p_value, phyper(q=lin_35_only_H3K9me2[i,4] -1, m=lin_35_only_H3K9me2[i,2], n=lin_35_only_H3K9me2[i,1]-lin_35_only_H3K9me2[i,2], k=lin_35_only_H3K9me2[i,3], lower.tail=FALSE))
}

lin_35_only_HT_adj_p_value <- p.adjust(lin_35_only_HT_p_value, method="fdr")
barplot_data[,3] = c(lin_35_only_H3K9me2$direct_targets_with_low_H3K9me2, lin_35_only_H3K9me2$n_direct_targets[1]-sum(lin_35_only_H3K9me2$direct_targets_with_low_H3K9me2))
write.table(cbind(row.names(lin_35_only_H3K9me2), lin_35_only_HT_FC, lin_35_only_HT_adj_p_value), sep="\t", file = paste(main_working_dir, "/H3K9me2/LIN35_only.lin35.hypergeom_test_results", sep=""), quote = F, row.names = F, col.names=F)

lin_35_only_H3K9me2_by_factor=data.frame(row.names=c("lin35", "lin36", "lin15B"), "tested_peaks" = rep(lin_35_only_H3K9me2[1,1], 3), "low_H3K9me2_in_mutant" = c(lin_35_only_H3K9me2[1,2]+lin_35_only_H3K9me2[4,2]+lin_35_only_H3K9me2[5,2]+lin_35_only_H3K9me2[7,2],
                                                                                                                                                                       lin_35_only_H3K9me2[2,2]+lin_35_only_H3K9me2[4,2]+lin_35_only_H3K9me2[6,2]+lin_35_only_H3K9me2[7,2],
                                                                                                                                                                       lin_35_only_H3K9me2[3,2]+lin_35_only_H3K9me2[5,2]+lin_35_only_H3K9me2[6,2]+lin_35_only_H3K9me2[7,2]),
                                            "n_direct_targets" = rep(lin_35_only_H3K9me2[1,3], 3), "direct_targets_with_low_H3K9me2" = c(lin_35_only_H3K9me2[1,4]+lin_35_only_H3K9me2[4,4]+lin_35_only_H3K9me2[5,4]+lin_35_only_H3K9me2[7,4],
                                                                                                                                            lin_35_only_H3K9me2[2,4]+lin_35_only_H3K9me2[4,4]+lin_35_only_H3K9me2[6,4]+lin_35_only_H3K9me2[7,4],
                                                                                                                                            lin_35_only_H3K9me2[3,4]+lin_35_only_H3K9me2[5,4]+lin_35_only_H3K9me2[6,4]+lin_35_only_H3K9me2[7,4]))
lin_35_only_by_factor_HT_FC=c()
lin_35_only_by_factor_HT_p_value=c()

for (i in c(1:dim(lin_35_only_H3K9me2_by_factor)[1])){
  lin_35_only_by_factor_HT_FC <-  append(lin_35_only_by_factor_HT_FC, (lin_35_only_H3K9me2_by_factor[i,4]/lin_35_only_H3K9me2_by_factor[i,3]) / (lin_35_only_H3K9me2_by_factor[i,2]/lin_35_only_H3K9me2_by_factor[i,1])) #  (x / k ) / (m / N)
  lin_35_only_by_factor_HT_p_value <- append(lin_35_only_by_factor_HT_p_value, phyper(q=lin_35_only_H3K9me2_by_factor[i,4] -1, m=lin_35_only_H3K9me2_by_factor[i,2], n=lin_35_only_H3K9me2_by_factor[i,1]-lin_35_only_H3K9me2_by_factor[i,2], k=lin_35_only_H3K9me2_by_factor[i,3], lower.tail=FALSE))
}
lin_35_only_by_factor_HT_adj_p_value=p.adjust(lin_35_only_by_factor_HT_p_value, method="fdr")


current_date=gsub("-", "", scan(text = as.character(Sys.time()), what = "")[1])
current_time=gsub(":", "", scan(text = as.character(Sys.time()), what = "")[2])

pdf(paste(main_working_dir, "/plots/H3K9me2_loss_direct_target_promoters_by_factor.", current_date, ".", current_time, ".pdf", sep=""), width=7, height=10)
par(mar=c(4,5,3,3), oma=c(0,0,0,8), mfrow=c(1,1))
barplot_color=c(t_col("coral2", 10), t_col("lightgreen", 10), t_col("mediumpurple1", 10))
k=barplot(matrix(c(lin_36_shared_H3K9me2_by_factor[,4]/lin_36_shared_H3K9me2_by_factor[,3],
                   lin_15B_shared_H3K9me2_by_factor[,4]/lin_15B_shared_H3K9me2_by_factor[,3]), ncol=2), ylim=c(0, 1), col=barplot_color, beside=T, las=1, 
          names.arg=c("LIN-36-shared", "LIN-15B-shared"), ylab="fraction of direct target promoters", main="H3K9me2 loss direct target promoters")
text(k, matrix(c(lin_36_shared_H3K9me2_by_factor[,4]/lin_36_shared_H3K9me2_by_factor[,3],
                 lin_15B_shared_H3K9me2_by_factor[,4]/lin_15B_shared_H3K9me2_by_factor[,3]), ncol=2), 
     labels = graphical_pvalue(c(lin_36_shared_by_factor_HT_adj_p_value, lin_15B_shared_by_factor_HT_adj_p_value)), pos=3)
text(k, matrix(c(lin_36_shared_H3K9me2_by_factor[,4]/lin_36_shared_H3K9me2_by_factor[,3],
                 lin_15B_shared_H3K9me2_by_factor[,4]/lin_15B_shared_H3K9me2_by_factor[,3]), ncol=2), 
     labels = c(lin_36_shared_H3K9me2_by_factor[,4], lin_15B_shared_H3K9me2_by_factor[,4]), pos=1)
par(fig = c(0, 1, 0, 1), oma = c(0, 0, 0, 0), mar = c(0, 0, 0, 0), new = TRUE)
plot(0, 0, type = "n", bty = "n", xaxt = "n", yaxt = "n")
legend(0.5, 0.5, legend = c("lin-35", "lin-36", "lin-15B"), fill = barplot_color, bty="n", ncol = 1, yjust = 1.5)
dev.off()
```
