## Supplementary material for "DREAM represses distinct targets by cooperating with different THAP domain proteins": Code and related files: HTZ1_90pct_enriched_genes.R

```
# extract top 10% of genes based on HTZ-1 coverage ofver the gene body (+500bp from Wormbase TSS to TTS)
args <- commandArgs()

main_working_dir = args[6]

HTZ1_wt_rep1 = read.table(paste(main_working_dir, "/HTZ1/elegans.gene_annotation.coding_gene_longest_isoform.500bp_gene_body.ce11.HTZ1_rep1_cov.bed", sep=""))
HTZ1_wt_rep2 = read.table(paste(main_working_dir, "/HTZ1/elegans.gene_annotation.coding_gene_longest_isoform.500bp_gene_body.ce11.HTZ1_rep2_cov.bed", sep=""))

HTZ1_wt = cbind(HTZ1_wt_rep1, HTZ1_wt_rep2$V5)
HTZ1_wt$HTZ1_mean=((HTZ1_wt[,5]+HTZ1_wt[,6])/2)/(HTZ1_wt$V3-HTZ1_wt$V2)

HTZ1_wt_filtered_10pct=HTZ1_wt[HTZ1_wt$HTZ1_mean > quantile(HTZ1_wt$HTZ1_mean, prob=0.90),]
write.table(HTZ1_wt_filtered_10pct[,c(1:4)], file = paste(main_working_dir, "/HTZ1/elegans.gene_annotation.coding_gene_longest_isoform.500bp_gene_body.ce11.HTZ1_90pct.bed", sep=""), sep = "\t", quote = F, col.names = F, row.names=F)
```
