## Supplementary material for "DREAM represses distinct targets by cooperating with different THAP domain proteins": Code and related files: HTZ1_direct_target_coverage.R

```
library("beeswarm")

args <- commandArgs()
main_working_dir = args[6]

t_col <- function(color, percent = 50, name = NULL) {
  rgb.val <- col2rgb(color)
  t.col <- rgb(rgb.val[1], rgb.val[2], rgb.val[3],
               max = 255,
               alpha = (100-percent)*255/100,
               names = name)
  invisible(t.col)
}

graphical_pvalue = function(x){
  x[x < 0.001] = "***"
  x[x > 0.001 & x < 0.01] = "**"
  x[x > 0.01 & x < 0.05] = "*"
  x[x >= 0.05] = "ns"
  print(x)
}

lin35_col=t_col("coral2", 10)
lin36full_col=t_col("lightgreen", 10)
lin36PM_col=t_col("mediumseagreen", 10)
lin15B_col=t_col("mediumpurple1", 10)

lin35_only_col=t_col("coral2", 50)
lin36_shared_col=t_col("lightgreen", 50)
lin15B_shared_col=t_col("mediumpurple1", 50)

# number of direct targets with high vs low gbHTZ-1 levels in N2
HTZ1_high_low_targets=read.table(paste(main_working_dir, "/HTZ1/high_low_gbHTZ1_direct_targets.txt", sep=""))
fisher.test(HTZ1_high_low_targets[c(1,2),])
fisher.test(HTZ1_high_low_targets[c(1,3),])
fisher.test(HTZ1_high_low_targets[c(2,3),])

current_date=gsub("-", "", scan(text = as.character(Sys.time()), what = "")[1])
current_time=gsub(":", "", scan(text = as.character(Sys.time()), what = "")[2])

pdf(paste(main_working_dir, "/plots/high_low_gbHTZ1_direct_targets.", current_date, ".", current_time, ".pdf", sep=""), width=6, height=6)
par(mfrow=c(1,1), mar=c(3,5,3,3), oma=c(2,0,0,0))
barplot(t(HTZ1_high_low_targets/rowSums(HTZ1_high_low_targets)), beside=F, las=1, names.arg = c("LIN-35 only", "LIN-36-shared", "LIN-15B-shared"), ylab="fraction of direct targets", main = "high gbHTZ-1 genes")
par(fig = c(0, 1, 0, 1), oma = c(0, 0, 0, 0), mar = c(0, 0, 0, 0), new = TRUE)
plot(0, 0, type = "n", bty = "n", xaxt = "n", yaxt = "n")
legend(0, -0.95, legend = c("high gbHTZ-1", "low gbHTZ-1"), fill = c("grey30", "grey90"), bty="n", ncol = 4, xjust = 0.2)
dev.off()

# number of genes and direct targets with lower gbHTZ-1 levels in mutants
HTZ1_downreg_genes_mutants=read.table(paste(main_working_dir, "/HTZ1/HTZ1_downreg_genes_mutants.txt", sep=""))
HTZ1_downreg_genes_mutants_matrix = cbind(HTZ1_downreg_genes_mutants$direct_targets, HTZ1_downreg_genes_mutants$all_downreg_HTZ1-HTZ1_downreg_genes_mutants$direct_targets)
fisher.test(HTZ1_downreg_genes_mutants_matrix)

pdf(paste(main_working_dir, "/plots/HTZ1_downreg_genes_mutants.", current_date, ".", current_time, ".pdf", sep=""), width=6, height=6)
par(mfrow=c(1,1), mar=c(3,5,3,3), oma=c(2,0,0,0))
barplot(t(HTZ1_downreg_genes_mutants_matrix), beside=F, las=1, names.arg = c(expression(italic("lin-35")), expression(italic("lin-36")), expression(italic("lin-15B"))), ylab="downreg gbHTZ-1", main = "gbHTZ-1 downregulated genes")
par(fig = c(0, 1, 0, 1), oma = c(0, 0, 0, 0), mar = c(0, 0, 0, 0), new = TRUE)
plot(0, 0, type = "n", bty = "n", xaxt = "n", yaxt = "n")
legend(0, -0.95, legend = c("direct targets", "not direct"), fill = c("grey30", "grey90"), bty="n", ncol = 4, xjust = 0.2)
dev.off()

direct_targets_with_HTZ1_downreg=read.table(paste(main_working_dir, "/HTZ1/direct_targets_with_HTZ1_downreg.txt", sep=""))
direct_targets_with_HTZ1_downreg = cbind(direct_targets_with_HTZ1_downreg$downreg_HTZ1, direct_targets_with_HTZ1_downreg$all_direct_targets-direct_targets_with_HTZ1_downreg$downreg_HTZ1)

pdf(paste(main_working_dir, "/plots/direct_targets_with_HTZ1_downreg.", current_date, ".", current_time, ".pdf", sep=""), width=6, height=6)

par(mfrow=c(1,1), mar=c(3,5,3,3), oma=c(2,0,0,0))
barplot(t(direct_targets_with_HTZ1_downreg), beside=F, las=1, names.arg = c(expression(italic("lin-35")), expression(italic("lin-36")), expression(italic("lin-15B"))), ylab="direct targets", main = "direct targets")
par(fig = c(0, 1, 0, 1), oma = c(0, 0, 0, 0), mar = c(0, 0, 0, 0), new = TRUE)
plot(0, 0, type = "n", bty = "n", xaxt = "n", yaxt = "n")
legend(0, -0.95, legend = c("HTZ-1 downreg", "not downreg"), fill = c("grey30", "grey90"), bty="n", ncol = 4, xjust = 0.2)
dev.off()

# compare HTZ-1 coverage over gene body of LIN15B-shared and LIN36-shared direct targets
lin15B_shared_HTZ1_N2 = read.table(paste(main_working_dir, "/HTZ1/genes_upregulated_lin15B_shared_vs_N2.direct.HTZ1_N2.bed", sep=""))
lin36_shared_HTZ1_N2 = read.table(paste(main_working_dir, "/HTZ1/genes_upregulated_lin36full_shared_vs_N2.direct.HTZ1_N2.bed", sep=""))
#lin35_only_HTZ1_N2 = read.table(paste(main_working_dir, "/HTZ1/genes_upregulated_lin35_only_vs_N2.direct.HTZ1_N2.bed", sep=""))

lin15B_lin36_P= wilcox.test(lin36_shared_HTZ1_N2$V4, lin15B_shared_HTZ1_N2$V4)$p.value

pdf(paste(main_working_dir, "/plots/HTZ1_direct_targets_coverage.N2.", current_date, ".", current_time, ".pdf", sep=""), width=6, height=6)
par(mfrow=c(1,1), mar=c(5,5,3,3))
x<-boxplot(lin36_shared_HTZ1_N2$V4,
           lin15B_shared_HTZ1_N2$V4, plot=F)
ylim_plot=c(min(c(x$stats, x$out)), max(c(x$stats, x$out)))
x<-boxplot(lin36_shared_HTZ1_N2$V4,lin15B_shared_HTZ1_N2$V4,notch=T, las=1, xaxt="n", ylim=c(ylim_plot[1], ylim_plot[2]),
           ylab="gbHTZ-1 coverage", main="direct targets gbHTZ-1 coverage", col=c(lin36_shared_col, lin15B_shared_col), outline=F)
text(cex=1, x=c(1,2), y=ylim_plot[1]-(ylim_plot[2]-ylim_plot[1])/15, c("LIN-36-shared", "LIN-15B-shared"), xpd=TRUE, srt=30, adj = 1)
beeswarm(list(lin36_shared_HTZ1_N2$V4,
              lin15B_shared_HTZ1_N2$V4), add=T, at = c(1,2),
         vertical = T, method = "swarm", spacing=1, corral="wrap", pch = 16, col = t_col("grey60", 50), cex = 0.6)
text(cex=1, x=1.5, y=ylim_plot[2]-(ylim_plot[2]-ylim_plot[1])/10, graphical_pvalue(lin15B_lin36_P), adj = c(0, 0))
segments(1, ylim_plot[2]-(ylim_plot[2]-ylim_plot[1])/10,
        2, ylim_plot[2]-(ylim_plot[2]-ylim_plot[1])/10)
dev.off()
```
