## Supplementary material for "DREAM represses distinct targets by cooperating with different THAP domain proteins": Code and related files: hypergeom_test_DE_gene_overlap.R

```
# hypergeometric test: LIN15B/LIN36 to LIN35 overlap
args <- commandArgs()

main_working_dir = args[6]

lin35_up = dim(read.table(paste(main_working_dir, "/DE_genes/direct_targets_all_rep/genes_upregulated_lin35_vs_N2.direct.txt", sep="")))[1] # number of target genes (m)
lin35_deseq = read.table(paste(main_working_dir, "/DE_genes/genes_DESeq_lin35_vs_N2.txt", sep=""))
lin35_deseq_exp = lin35_deseq[!is.na(lin35_deseq$padj),]


lin15B_up = dim(read.table(paste(main_working_dir, "/DE_genes/direct_targets_all_rep/genes_upregulated_lin15B_vs_N2.direct.txt", sep="")))[1] # size of selection (k)
lin15B_deseq = read.table(paste(main_working_dir, "/DE_genes/genes_DESeq_lin15B_vs_N2.txt", sep=""))
lin15B_deseq_exp = lin15B_deseq[!is.na(lin15B_deseq$padj),]

lin15B_lin35_exp = dim(lin15B_deseq_exp[row.names(lin15B_deseq_exp) %in% row.names(lin35_deseq_exp),])[1] # size of universe (N)
non_lin35_up = lin15B_lin35_exp-lin35_up # number of non-target genes (n)

lin15B_lin35_overlap = dim(read.table(paste(main_working_dir, "/DE_genes/direct_targets_all_rep/genes_upregulated_lin15B_lin35.direct.txt", sep="")))[1] # number of targets in selection (x)

fold.enrichment <-  (lin15B_lin35_overlap/lin15B_up) / (lin35_up/lin15B_lin35_exp) #  (x / k ) / (m / N)
p.value <-  phyper(q=lin15B_lin35_overlap -1, m=lin35_up, n=non_lin35_up, k=lin15B_up, lower.tail=FALSE)

cat(paste("LIN-15B+LIN-35: fold enrichment:", fold.enrichment, "; p.value:", p.value, "\n"))

lin36full_up = dim(read.table(paste(main_working_dir, "/DE_genes/direct_targets_all_rep/genes_upregulated_lin36full_vs_N2.direct.txt", sep="")))[1] # size of selection (k)
lin36full_deseq = read.table(paste(main_working_dir, "/DE_genes/genes_DESeq_lin36full_vs_N2.txt", sep=""))
lin36full_deseq_exp = lin36full_deseq[!is.na(lin36full_deseq$padj),]

lin36full_lin35_exp = dim(lin36full_deseq_exp[row.names(lin36full_deseq_exp) %in% row.names(lin35_deseq_exp),])[1] # size of universe (N)
non_lin35_up = lin36full_lin35_exp-lin35_up # number of non-target genes (n)

lin36full_lin35_overlap = dim(read.table(paste(main_working_dir, "/DE_genes/direct_targets_all_rep/genes_upregulated_lin36_lin35.direct.txt", sep="")))[1] # number of targets in selection (x)

fold.enrichment <-  (lin36full_lin35_overlap/lin36full_up) / (lin35_up/lin36full_lin35_exp) #  (x / k ) / (m / N)
p.value <-  phyper(q=lin36full_lin35_overlap -1, m=lin35_up, n=non_lin35_up, k=lin36full_up, lower.tail=FALSE)

cat(paste("LIN-36+LIN-35: fold enrichment:", fold.enrichment, "; p.value:", p.value, "\n"))
```
