## Supplementary material for "DREAM represses distinct targets by cooperating with different THAP domain proteins": Code and related files: LIN15BTHAPdel_expression_analysis.R

```
args <- commandArgs()
main_working_dir = args[6]

t_col <- function(color, percent = 50, name = NULL) {
  rgb.val <- col2rgb(color)
  t.col <- rgb(rgb.val[1], rgb.val[2], rgb.val[3],
               max = 255,
               alpha = (100-percent)*255/100,
               names = name)
  invisible(t.col)
}

graphical_pvalue = function(x){
  x[x < 0.001] = "***"
  x[x > 0.001 & x < 0.01] = "**"
  x[x > 0.01 & x < 0.05] = "*"
  x[x >= 0.05] = "ns"
  print(x)
}

lin15B_DESeq = read.table(paste(main_working_dir, "/DE_genes/genes_DESeq_lin15BTHAPdel_vs_N2.txt", sep=""))
DESeq_coverage = read.table(paste(main_working_dir, "/DE_genes/normalized_counts_DESeq.txt", sep=""))

high_LIN15BTHAPdel_genes=read.table(paste(main_working_dir, "/THAPdel_mutants/genes_stronger_LIN15BTHAPdel_peaks.genes", sep=""))
control_LIN15BTHAPdel_genes=read.table(paste(main_working_dir, "/THAPdel_mutants/genes_control_LIN15BTHAPdel_peaks.genes", sep=""))

lfc_P = wilcox.test(lin15B_DESeq$log2FoldChange[row.names(lin15B_DESeq) %in% high_LIN15BTHAPdel_genes$V1], lin15B_DESeq$log2FoldChange[row.names(lin15B_DESeq) %in% control_LIN15BTHAPdel_genes$V1])$p.value


current_date=gsub("-", "", scan(text = as.character(Sys.time()), what = "")[1])
current_time=gsub(":", "", scan(text = as.character(Sys.time()), what = "")[2])

pdf(paste(main_working_dir, "/plots/lin15BTHAPdel_gene_expression.", current_date, ".", current_time, ".pdf", sep=""), width=4.5, height=5)

par(mfrow=c(1,1), mar=c(4,5,3,3))
x<-boxplot(lin15B_DESeq$log2FoldChange[row.names(lin15B_DESeq) %in% high_LIN15BTHAPdel_genes$V1],
	lin15B_DESeq$log2FoldChange[row.names(lin15B_DESeq) %in% control_LIN15BTHAPdel_genes$V1], plot=F)
ylim_box=c(min(c(x$stats, x$out)), max(c(x$stats, x$out)))
x<-boxplot(lin15B_DESeq$log2FoldChange[row.names(lin15B_DESeq) %in% high_LIN15BTHAPdel_genes$V1], lin15B_DESeq$log2FoldChange[row.names(lin15B_DESeq) %in% control_LIN15BTHAPdel_genes$V1], notch=T,
           main="expression of LIN-15B-associated genes in lin-15B(THAPdel)", ylab="LFC expression lin-15B(THAPdel) vs N2", las=1,
           ylim=c(ylim_box[1], ylim_box[2]+(ylim_box[2]-ylim_box[1])/15), names=c("stronger LIN-15B signal", "control"), outline=F)
abline(h=0, lty=2, lwd=2, col="grey")
text(1.5, ylim_box[2]-((ylim_box[2]-ylim_box[1])/15), labels = graphical_pvalue(lfc_P), pos = 3)

dev.off()
```
