## Supplementary material for "DREAM represses distinct targets by cooperating with different THAP domain proteins": Code and related files: met2_analysis.R

```
library("beeswarm")

args <- commandArgs()
main_working_dir = args[6]

t_col <- function(color, percent = 50, name = NULL) {
  rgb.val <- col2rgb(color)
  t.col <- rgb(rgb.val[1], rgb.val[2], rgb.val[3],
               max = 255,
               alpha = (100-percent)*255/100,
               names = name)
  invisible(t.col)
}

graphical_pvalue = function(x){
  x[x < 0.001] = "***"
  x[x > 0.001 & x < 0.01] = "**"
  x[x > 0.01 & x < 0.05] = "*"
  x[x >= 0.05] = "ns"
  print(x)
}

met2_DESeq = read.table(paste(main_working_dir, "/DE_genes/genes_DESeq_met2_vs_N2.txt", sep=""))
DESeq_coverage = read.table(paste(main_working_dir, "/DE_genes/normalized_counts_DESeq.txt", sep=""))

LIN35_specific=read.table(paste(main_working_dir, "/DE_genes/direct_targets_all_rep/genes_upregulated_lin35_specific.direct.txt", sep=""))
LIN36_shared=read.table(paste(main_working_dir, "/DE_genes/direct_targets_all_rep/genes_upregulated_lin36_shared.direct.txt", sep=""))
LIN15B_shared=read.table(paste(main_working_dir, "/DE_genes/direct_targets_all_rep/genes_upregulated_lin15B_shared.direct.txt", sep=""))

lin35_only_col=t_col("coral2", 50)
lin36_shared_col=t_col("lightgreen", 50)
lin15B_shared_col=t_col("mediumpurple1", 50)

LIN15B_36_met2 = matrix(c(dim(met2_DESeq[met2_DESeq$log2FoldChange > 0 & met2_DESeq$padj < 0.01 & row.names(met2_DESeq) %in% LIN36_shared$V1,])[1],
                          dim(LIN36_shared)[1] - dim(met2_DESeq[met2_DESeq$log2FoldChange > 0 & met2_DESeq$padj < 0.01 & row.names(met2_DESeq) %in% LIN36_shared$V1,])[1],
                          dim(met2_DESeq[met2_DESeq$log2FoldChange > 0 & met2_DESeq$padj < 0.01 & row.names(met2_DESeq) %in% LIN15B_shared$V1,])[1],
                          dim(LIN15B_shared)[1] - dim(met2_DESeq[met2_DESeq$log2FoldChange > 0 & met2_DESeq$padj < 0.01 & row.names(met2_DESeq) %in% LIN15B_shared$V1,])[1]), ncol=2, byrow=T)
fisher.test(LIN15B_36_met2)

LIN36_shared_exp = DESeq_coverage[rownames(DESeq_coverage) %in% LIN36_shared$V1, c(30, 31, 32, 26, 27)]
LIN15B_shared_exp = DESeq_coverage[rownames(DESeq_coverage) %in% LIN15B_shared$V1, c(30, 31, 32, 26, 27)]

LIN36_shared_exp$N2_avg = (LIN36_shared_exp[,1] + LIN36_shared_exp[,2] + LIN36_shared_exp[,3])/3
LIN36_shared_exp$met2_avg = (LIN36_shared_exp[,4] + LIN36_shared_exp[,5])/2

LIN15B_shared_exp$N2_avg = (LIN15B_shared_exp[,1] + LIN15B_shared_exp[,2] + LIN15B_shared_exp[,3])/3
LIN15B_shared_exp$met2_avg = (LIN15B_shared_exp[,4] + LIN15B_shared_exp[,5])/2

lfc_P = c()
lfc_P = append(lfc_P, wilcox.test(LIN36_shared_exp$N2_avg, LIN36_shared_exp$met2_avg, paired=T)$p.value)
lfc_P = append(lfc_P, wilcox.test(LIN15B_shared_exp$N2_avg, LIN15B_shared_exp$met2_avg, paired=T)$p.value)
lfc_P_adj=p.adjust(lfc_P, method="fdr")

current_date=gsub("-", "", scan(text = as.character(Sys.time()), what = "")[1])
current_time=gsub(":", "", scan(text = as.character(Sys.time()), what = "")[2])


pdf(paste(main_working_dir, "/plots/met2_gene_expression.", current_date, ".", current_time, ".pdf", sep=""), width=4.5, height=5)

par(mfrow=c(1,1), mar=c(4,5,3,3))
x<-boxplot(met2_DESeq$log2FoldChange[row.names(met2_DESeq) %in% LIN36_shared$V1],
           met2_DESeq$log2FoldChange[row.names(met2_DESeq) %in% LIN15B_shared$V1], plot=F)
ylim_box=c(min(c(x$stats, x$out)), max(c(x$stats, x$out)))
x<-boxplot(met2_DESeq$log2FoldChange[row.names(met2_DESeq) %in% LIN36_shared$V1],
           met2_DESeq$log2FoldChange[row.names(met2_DESeq) %in% LIN15B_shared$V1], notch=T, 
           main="direct targets expression in met-2", ylab="LFC expression met-2 vs N2", las=1,
           col=c(lin36_shared_col, lin15B_shared_col), ylim=c(ylim_box[1], ylim_box[2]+(ylim_box[2]-ylim_box[1])/15), names=c("LIN-36-shared", "LIN-15B-shared"), outline=F)
abline(h=0, lty=2, lwd=2, col="grey")
beeswarm(list(met2_DESeq$log2FoldChange[row.names(met2_DESeq) %in% LIN36_shared$V1],
              met2_DESeq$log2FoldChange[row.names(met2_DESeq) %in% LIN15B_shared$V1]), add=T, at = c(1,2),
         vertical = T, method = "swarm", spacing=1, corral="wrap", pch = 16, col = t_col("grey60", 50), cex = 0.6)
text(c(1, 2), ylim_box[2]-((ylim_box[2]-ylim_box[1])/15), labels = graphical_pvalue(lfc_P_adj), pos = 3)


dev.off()
```
