## Supplementary material for "DREAM represses distinct targets by cooperating with different THAP domain proteins": Code and related files: motif_enrichment_launcher.sh

```
#!/bin/bash
#
#SBATCH --job-name=motiffind
#SBATCH --output=/mnt/home1/ahringer/fnc21/slurm_out/motif.%N.%j.out
#SBATCH --error=/mnt/home1/ahringer/fnc21/slurm_err/motif.%N.%j.err
#SBATCH --ntasks=1
#SBATCH -N 1
#SBATCH -n 10
#SBATCH --mem-per-cpu=4000
#SBATCH --mail-type=END,FAIL
#SBATCH --mail-user=

output_dir=$1
factor=$2
control_group=$3
peak_set=$4
control_peaks=$5

output_dir_meme=$output_dir/meme

mkdir -p $output_dir_meme

if [ $control_group == "scrambled" ]
then
    meme -oc $output_dir_meme/$factor.$control_group -objfun de -dna -revcomp -nmotifs 10 -p 10 $peak_set
else
    meme -oc $output_dir_meme/$factor.$control_group -objfun de -neg $control_peaks -dna -revcomp -nmotifs 10 -p 10 $peak_set 
fi
```
