## Supplementary material for "DREAM represses distinct targets by cooperating with different THAP domain proteins": Code and related files: other_DREAM_upregulation.R

```
args <- commandArgs()
main_working_dir = args[6]

t_col <- function(color, percent = 50, name = NULL) {
  rgb.val <- col2rgb(color)
  t.col <- rgb(rgb.val[1], rgb.val[2], rgb.val[3],
               max = 255,
               alpha = (100-percent)*255/100,
               names = name)
  invisible(t.col)
  
}

graphical_pvalue = function(x){
  x[x < 0.001] = "***"
  x[x > 0.001 & x < 0.01] = "**"
  x[x > 0.01 & x < 0.05] = "*"
  x[x >= 0.05] = "ns"
  print(x)
}

barplot_data=matrix(rep(0, 6), ncol=1)
barplot_p=matrix(rep(0, 6), ncol=1)

DREAM_DR = read.table(paste(main_working_dir, "/other_DREAM_factors/other_DREAM.hypergeom_overlap", sep=""))
DREAM_DR_HT_FC=c()
DREAM_DR_HT_p_value=c()

for (i in c(1:dim(DREAM_DR)[1])){
  DREAM_DR_HT_FC <-  append(DREAM_DR_HT_FC, (DREAM_DR[i,4]/DREAM_DR[i,3]) / (DREAM_DR[i,2]/DREAM_DR[i,1])) #  (x / k ) / (m / N)
  DREAM_DR_HT_p_value <- append(DREAM_DR_HT_p_value, phyper(q=DREAM_DR[i,4] -1, m=DREAM_DR[i,2], n=DREAM_DR[i,1]-DREAM_DR[i,2], k=DREAM_DR[i,3], lower.tail=FALSE))
}

DREAM_DR_fisher_p_value=c()
DREAM_DR_fisher_p_value=append(DREAM_DR_fisher_p_value, fisher.test(matrix(c(DREAM_DR[1,4], DREAM_DR[1,3]-DREAM_DR[1,4], DREAM_DR[2,4], DREAM_DR[2,3]-DREAM_DR[2,4]), ncol=2, byrow=T))$p.value)
DREAM_DR_fisher_p_value=append(DREAM_DR_fisher_p_value, fisher.test(matrix(c(DREAM_DR[3,4], DREAM_DR[3,3]-DREAM_DR[3,4], DREAM_DR[4,4], DREAM_DR[4,3]-DREAM_DR[4,4]), ncol=2, byrow=T))$p.value)
DREAM_DR_fisher_p_value=append(DREAM_DR_fisher_p_value, fisher.test(matrix(c(DREAM_DR[5,4], DREAM_DR[5,3]-DREAM_DR[5,4], DREAM_DR[6,4], DREAM_DR[6,3]-DREAM_DR[6,4]), ncol=2, byrow=T))$p.value)

DREAM_DR_HT_adj_p_value <- p.adjust(DREAM_DR_HT_p_value, method="fdr")
barplot_data[,1] = c(DREAM_DR$direct_targets_upreg_in_mutant/DREAM_DR$n_direct_targets)
barplot_p[,1]=DREAM_DR_HT_adj_p_value
write.table(cbind(row.names(DREAM_DR), DREAM_DR_HT_FC, DREAM_DR_HT_adj_p_value), sep="\t", file = paste(main_working_dir, "/other_DREAM_factors/other_DREAM.hypergeom_results", sep=""), quote = F, row.names = F, col.names=F)

current_date=gsub("-", "", scan(text = as.character(Sys.time()), what = "")[1])
current_time=gsub(":", "", scan(text = as.character(Sys.time()), what = "")[2])

pdf(paste(main_working_dir, "/plots/shared_direct_targets_upregulation_other_DREAM_mutants.", current_date, ".", current_time, ".pdf", sep=""), width=8, height=8)
par(mar=c(4,5,3,3), oma=c(0,0,0,0), mfrow=c(1,1))
k=barplot(rep(NA, 3), ylim=c(-1,1), axes=F) 
abline(h=c(0, 0.5, -0.5), lty=2, lwd=2, col=c("grey", t_col("indianred", 50), t_col("indianred", 50)), )
k=barplot(height = barplot_data[c(1,3,5),1], axes=F, beside=T, las=1, add=T, main = "shared direct targets upregulated in other DREAM mutants", col=t_col("lightgreen", 0)) 
k=barplot(height = -barplot_data[c(2,4,6),1], axes=F, beside=T, las=1, add=T, col=t_col("mediumpurple1", 0)) 
axis(2, at = c(-1, -0.5, 0, 0.5, 1), labels = c(1, 0.5, 0, 0.5, 1), las=1)
text(-0.3, 0.5, labels = "LIN36-shared upregulated dt", srt = 90, xpd=T, cex=0.9)
text(-0.3, -0.5, labels = "LIN15B-shared upregulated dt", srt = 90, xpd=T, cex=0.9)
text(k[,1], 1, graphical_pvalue(DREAM_DR_fisher_p_value), pos = 1)
text(k[,1], -1.15, labels = rep(c("dpl-1", "efl-1", "lin-37"), 2), xpd=T)
dev.off()
```
