## Supplementary material for "DREAM represses distinct targets by cooperating with different THAP domain proteins": Code and related files: star_rnaseq_analysis.sh

```
#!/bin/bash
#
#SBATCH --job-name=star
#SBATCH --output=/mnt/home1/ahringer/fnc21/slurm_out/star.%N.%j.out
#SBATCH --error=/mnt/home1/ahringer/fnc21/slurm_err/star.%N.%j.err
#SBATCH --ntasks=1
#SBATCH -N 1
#SBATCH -n 6
#SBATCH --mem-per-cpu=4000
#SBATCH --mail-type=END,FAIL
#SBATCH --mail-user=

working_dir=$1
input_file=$2
experiment=$3
species=$4
strain=$5

unstranded_tag="--outSAMstrandField intronMotif"
genome_idx=$working_dir/STAR_idx/$species
output_path_name=$working_dir/data/$species/$experiment/alignment/$strain
genome_path=$working_dir/species/$species/genome
fastq_path=$working_dir/data/$species/$experiment/fastq
genes_GTF_path=$working_dir/species/$species/gene_annotation/$species.annotations.coding_genes.ce11.gtf

mkdir -p $output_path_name

if [ ! -d "$genome_idx" ];
then
    mkdir -p $genome_idx
    STAR --genomeSAindexNbases 10 --sjdbGTFfile $genes_GTF_path --runMode genomeGenerate --runThreadN 8 --genomeDir $genome_idx --genomeFastaFiles $genome_path/$species.fa
fi

#~/software/STAR-2.5.4b/bin/Linux_x86_64/STAR --readFilesCommand zcat --runThreadN 6 --genomeDir $genome_idx --readFilesIn $fastq_path"/"$strain"."$input_file".fq.gz" --outFileNamePrefix $output_path_name"/"$strain"."$input_file"." --outSAMtype BAM SortedByCoordinate --outSAMattrIHstart 0 --outWigType wiggle --twopassMode Basic $unstranded_tag

~/software/wigToBigWig $output_path_name"/"$strain"."$input_file".Signal.UniqueMultiple.str1.out.wig" $genome_path/$species.chrom.sizes.txt $output_path_name"/"$strain"."$input_file".Signal.UniqueMultiple.str1.out.bw"
~/software/wigToBigWig $output_path_name"/"$strain"."$input_file".Signal.UniqueMultiple.str2.out.wig" $genome_path/$species.chrom.sizes.txt $output_path_name"/"$strain"."$input_file".Signal.UniqueMultiple.str2.out.bw"
```
