## Supplementary material for "DREAM represses distinct targets by cooperating with different THAP domain proteins": Code and related files: upregulated_genes_expression.R

```
# GL enrichment among upregulated genes in all mutants
args <- commandArgs()

main_working_dir = args[6]

gene_exp = read.table(paste(main_working_dir, "/species/elegans/Cao_single_cell_TPM_by_cell_type.txt", sep=""), header = T)
t_col <- function(color, percent = 50, name = NULL) {
  rgb.val <- col2rgb(color)
  t.col <- rgb(rgb.val[1], rgb.val[2], rgb.val[3],
               max = 255,
               alpha = (100-percent)*255/100,
               names = name)
  invisible(t.col)
  
}

graphical_pvalue = function(x){
  x[x < 0.001] = "***"
  x[x > 0.001 & x < 0.01] = "**"
  x[x > 0.01 & x < 0.05] = "*"
  x[x >= 0.05] = "ns"
  print(x)
}

lin35_col=t_col("coral2", 10)
lin36full_col=t_col("lightgreen", 10)
lin15B_col=t_col("mediumpurple1", 10)

lin35_only_col=t_col("coral2", 50)
lin36_shared_col=t_col("lightgreen", 50)
lin15B_shared_col=t_col("mediumpurple1", 50)

cell_types=names(gene_exp)[c(3:dim(gene_exp)[2])]

lin15B_exp=read.table(paste(main_working_dir, "/DE_genes/gene_expression/genes_upregulated_lin15B_vs_N2.expression", sep=""), col.names=names(gene_exp))[,c(3:29)]
lin35_exp=read.table(paste(main_working_dir, "/DE_genes/gene_expression/genes_upregulated_lin35_vs_N2.expression", sep=""), col.names=names(gene_exp))[,c(3:29)]
lin36full_exp=read.table(paste(main_working_dir, "/DE_genes/gene_expression/genes_upregulated_lin36full_vs_N2.expression", sep=""), col.names=names(gene_exp))[,c(3:29)]

lin15B_exp_gl_enrichment = c()
lin35_exp_gl_enrichment = c()
lin36full_exp_gl_enrichment = c()

for (i in c(2:dim(lin15B_exp)[2])){
  lin15B_exp_gl_enrichment=append(lin15B_exp_gl_enrichment, wilcox.test(lin15B_exp[,1], lin15B_exp[,i])$p.value)
  lin35_exp_gl_enrichment=append(lin35_exp_gl_enrichment, wilcox.test(lin35_exp[,1], lin35_exp[,i])$p.value)
  lin36full_exp_gl_enrichment=append(lin36full_exp_gl_enrichment, wilcox.test(lin36full_exp[,1], lin36full_exp[,i])$p.value)
}

which(p.adjust(lin15B_exp_gl_enrichment, method = "fdr") > 0.01)
which(p.adjust(lin35_exp_gl_enrichment, method = "fdr") > 0.01)
which(p.adjust(lin36full_exp_gl_enrichment, method = "fdr") > 0.01)

current_date=gsub("-", "", scan(text = as.character(Sys.time()), what = "")[1])
current_time=gsub(":", "", scan(text = as.character(Sys.time()), what = "")[2])
pdf(paste(main_working_dir, "/plots/gonad_expression_upreg_genes.", current_date, ".", current_time, ".pdf", sep=""), width=14, height=6)

types_to_test=c("Germline", "Somatic_gonad_precursors", "Seam_cells", "Intestine", "Non.seam_hypodermis", "Body_wall_muscle")

col_to_test=c()
for (i in c(1:length(types_to_test))){
  col_to_test=append(col_to_test, which(match(cell_types, types_to_test)==i))
}


boxplot_at=c(1:6, 8:13, 15:20)

x = boxplot(c(log2(lin35_exp[,col_to_test]+1e-01),
              log2(lin36full_exp[,col_to_test]+1e-01),
              log2(lin15B_exp[,col_to_test]+1e-01)), plot=F)
ylim_plot=c(min(c(x$stats, x$out)), max(c(x$stats, x$out)))

par(mfrow=c(1,1), oma=c(2,0,0,0), mar=c(4,4,3,3), pch=21)
boxplot(c(log2(lin35_exp[,col_to_test]+1e-01),
          log2(lin36full_exp[,col_to_test]+1e-01),
          log2(lin15B_exp[,col_to_test]+1e-01)), las=1, ylab = "log2 TPM", notch=T, at = boxplot_at,
        col=rep(c("darkorchid2", "deepskyblue1", "deepskyblue3", "deepskyblue4", "darkorange1", "darkorange3"), length(types_to_test)), xaxt="n", main="gonad expression upregulated genes", outcol=rgb(0, 0, 0, .99))
abline(h=0, lty=2, col="grey")
axis(1, at = boxplot_at, label=NA)
text(cex=1, x=boxplot_at, y=ylim_plot[1]-(ylim_plot[2]-ylim_plot[1])/10, rep(c("GL", "SGP", "SC", "I", "NSH", "BWM"), 3), xpd=TRUE, srt=30, adj = 1)
par(fig = c(0, 1, 0, 1), oma = c(0, 0, 0, 0), mar = c(0, 0, 0, 0), new = TRUE)
plot(0, 0, type = "n", bty = "n", xaxt = "n", yaxt = "n")
legend(0, -0.9, legend = c("Germline", "dividing cell types", "non-dividing cell types"), fill = c("darkorchid2", "deepskyblue1", "darkorange1"), bty="n", ncol = 3, xjust = 0.5)
dev.off()

lin15B_shared=read.table(paste(main_working_dir, "/DE_genes/direct_targets_all_rep/genes_upregulated_lin15B_shared.direct.txt", sep=""))
lin36_shared=read.table(paste(main_working_dir, "/DE_genes/direct_targets_all_rep/genes_upregulated_lin36_shared.direct.txt", sep=""))
lin35_only=read.table(paste(main_working_dir, "/DE_genes/direct_targets_all_rep/genes_upregulated_lin35_specific.direct.txt", sep=""))

lin15B_exp=read.table(paste(main_working_dir, "/DE_genes/gene_expression/genes_upregulated_lin15B_vs_N2.expression", sep=""), col.names=names(gene_exp))
lin36full_exp=read.table(paste(main_working_dir, "/DE_genes/gene_expression/genes_upregulated_lin36full_vs_N2.expression", sep=""), col.names=names(gene_exp))
lin35_exp=read.table(paste(main_working_dir, "/DE_genes/gene_expression/genes_upregulated_lin35_vs_N2.expression", sep=""), col.names=names(gene_exp))

lin15B_shared_exp=lin15B_exp[lin15B_exp$gene_id %in% lin15B_shared$V1,c(3:dim(lin15B_exp)[2])]
lin36full_shared_exp=lin36full_exp[lin36full_exp$gene_id %in% lin36_shared$V1,c(3:dim(lin36full_exp)[2])]
lin35_only_exp=lin35_exp[lin35_exp$gene_id %in% lin35_only$V1,c(3:dim(lin35_exp)[2])]

lin15B_shared_GL_spec = lin15B_shared_exp[,1]/rowSums(lin15B_shared_exp[,c(2:dim(lin15B_shared_exp)[2])])
lin36full_shared_GL_spec = lin36full_shared_exp[,1]/rowSums(lin36full_shared_exp[,c(2:dim(lin36full_shared_exp)[2])])
lin35_only_GL_spec = lin35_only_exp[,1]/rowSums(lin35_only_exp[,c(2:dim(lin35_only_exp)[2])])

lin15B_shared_GL_spec = lin15B_shared_exp[,1]/rowSums(lin15B_shared_exp)
lin36full_shared_GL_spec = lin36full_shared_exp[,1]/rowSums(lin36full_shared_exp)
lin35_only_GL_spec = lin35_only_exp[,1]/rowSums(lin35_only_exp)

all_p = c()
all_p = append(all_p, wilcox.test(lin15B_shared_GL_spec, lin36full_shared_GL_spec)$p.value)
all_p = append(all_p, wilcox.test(lin15B_shared_GL_spec, lin35_only_GL_spec)$p.value)
all_p = append(all_p, wilcox.test(lin35_only_GL_spec, lin36full_shared_GL_spec)$p.value)

pdf(paste(main_working_dir, "/plots/germline_specificity_direct_targets.", current_date, ".", current_time, ".pdf", sep=""), width=6, height=6)
par(mfrow=c(1,1), oma=c(0,0,0,0), mar=c(4,5,3,3))
x = boxplot(lin35_only_GL_spec,
            lin36full_shared_GL_spec,
            lin15B_shared_GL_spec, plot=F)
ylim_plot=c(min(c(x$stats, x$out)), max(c(x$stats, x$out)))

boxplot(lin35_only_GL_spec,
        lin36full_shared_GL_spec,
        lin15B_shared_GL_spec, notch=T, main = "GL specificity direct targets", ylab = "log2 TPM", names=c("LIN-35 only", "LIN-36 shared", "LIN-15B shared"), 
        col=c(lin35_only_col, lin36_shared_col, lin15B_shared_col), ylim = c(ylim_plot[1], ylim_plot[2]+(ylim_plot[2]-ylim_plot[1])/8), las=1, outcol=rgb(0, 0, 0, .99))
text(cex=1, x=c(1.5, 2, 2.5), y=c(ylim_plot[2]+((ylim_plot[2]-ylim_plot[1])/25)*1,
                                  ylim_plot[2]+((ylim_plot[2]-ylim_plot[1])/25)*2,
                                  ylim_plot[2]+((ylim_plot[2]-ylim_plot[1])/25)*3), graphical_pvalue(all_p), adj = c(0, 0))
segments(c(1,1,2), c(ylim_plot[2]+((ylim_plot[2]-ylim_plot[1])/25)*1,
                     ylim_plot[2]+((ylim_plot[2]-ylim_plot[1])/25)*2,
                     ylim_plot[2]+((ylim_plot[2]-ylim_plot[1])/25)*3), 
         c(2,3,3), c(ylim_plot[2]+((ylim_plot[2]-ylim_plot[1])/25)*1,
                     ylim_plot[2]+((ylim_plot[2]-ylim_plot[1])/25)*2,
                     ylim_plot[2]+((ylim_plot[2]-ylim_plot[1])/25)*3))
dev.off()
```
